# Pharmacological inhibition of 11ßhydroxysteroid dehydrogenase type 1 after myocardial infarction targets extracellular matrix processing and preserves cardiac function in a translational mini-pig model

**DOI:** 10.1101/2024.12.02.626322

**Authors:** S Al Disi, R Ascione, S Khan, T Johnson, E Sammut, VD Bruno, DB Lopez, CA James, J Simpson, N Homer, M Millar, T Singh, A von Kreigsheim, NL Mills, BR Walker, R Andrew, SP Webster, A Whittaker, A Freeman, GA Gray

## Abstract

**Background and Purpose:** Plasma glucocorticoids (GCs) increase acutely after myocardial infarction (MI), thereafter tissue levels are amplified selectively within cells expressing 11ßHydroxysteroid Dehydrogenase type 1 (11ßHSD1) that regenerates active GCs from circulating metabolites. GCs initially protect cardiomyocytes and prevent excessive inflammation after MI but can also suppress subsequent wound repair leading to functional decline. The present study aimed to investigate the potential of pharmacological 11ßHSD1 inhibition after MI to prevent deterioration of cardiac function and its impact on wound repair.

**Experimental Approach:** Adult female Gottingen mini-pigs underwent percutaneous balloon MI/reperfusion and were randomised to receive either oral 11ßHSD1 inhibitor AZD8329 (n=11), or vehicle (n=9), from 2 until 27 days later, with concurrent administration of clinically relevant therapeutic intervention (anti-platelet, statin and ACE inhibitor).

**Key Results:** AZD8329 treatment increased plasma accumulation of cortisone substrate consistent with successful 11ßHSD1 inhibition. Gadolinium-enhanced MRI showed equivalent infarct size in both groups prior to commencing treatment. 28 days after MI cardiac function and LV area were preserved in the AZD8329 treated group relative to vehicle. There was no impact of 11ßHSD1 inhibitor on neovascularisation or infarct area. Mass spectrometry imaging revealed AZD8329 binding to the healing infarct and altered regulation of extracellular matrix (ECM) processing was highlighted by birefringence microscopy and proteomic analysis.

**Conclusions and Implications:** Pharmacological inhibition of 11ßHSD1 after MI prevents deterioration of cardiac function and detrimental remodelling. 11ßHSD1 inhibitors have safely reached phase 2 clinical trials in diabetes and dementia and could be repurposed as an addition to standard care after MI to prevent the development of heart failure.

**Bullet Point Summary:** *What is already known?:*

- GCs are released from the adrenal gland after MI, but also regenerated within the heart from circulating precursors by the enzyme 11ßHSD1 in cardiomyocytes, fibroblasts and macrophages.
- Genetic suppression of *Hsd11b1* expression in the mouse promotes neovascularisation, prevents infarct expansion during infarct repair after MI and the development of heart failure.

*What does this study add?:*

- Oral pharmacological inhibition of 11ßHSD1 after MI/reperfusion in a translational mini-pig model of MI receiving concurrent clinically relevant therapy prevents cardiac functional deterioration and adverse ventricular remodelling over the following 4 weeks.
- Mass spectrometry imaging reveals target engagement of the 11ßHSD1i in the repairing infarct.
- The mechanism is independent of neovascularisation but does involve modification of extracellular matrix remodelling during repair and scar formation.

*What is the clinical significance?:*

- Tissue 11ßHSD1 expression is increased in aging when the risk of MI is higher.
- Pharmacological inhibitors of 11ßHSD1 have safely reached phase 2 clinical trials for dementia and metabolic disease and could be repurposed for use post-MI to prevent the development of heart failure.

## 1. INTRODUCTION

Acute survival following myocardial infarction (MI) has dramatically increased thanks to improved early management (Byrne et al., 2023). However, this has resulted in many more individuals surviving with damage to their heart that increases the likelihood of subsequent adverse remodelling and the development of heart failure (HF) (BHF, 2024)). Recent innovations in drug intervention, particularly the metabolic and haemodynamic actions of <u>sodium/glucose co-transporter 2</u> (SGLT2) inhibitors and <u>Angiotensin Receptor</u> <u>Neprilysin</u> Inhibitors (ARNIs), have had significant positive impact in HF (Bozkurt, 2024; Lam, Docherty, Ho, McMurray, Myhre & Omland, 2023). In the case of HF following MI, a specific window of opportunity for intervention remains in the early post-MI period to optimise infarct repair and reduce the stimulus for the remodelling that leads to HF.

Glucocorticoids (GCs) released acutely in response to hypothalamo-pituitary-adrenal (HPA) axis activation after MI initially protect cardiomyocytes from ischaemic cell death (Libby, Maroko, Bloor, Sobel & Braunwald, 1973; Skyschally et al., 2004; Tokudome et al., 2009). GCs are also generated within cells expressing the enzyme <u>11ßHydroxysteroid Dehydrogenase</u> <u>type 1</u> (11ßHSD1) that regenerates active GCs from circulating metabolites (Chapman, Holmes & Seckl, 2013). In the heart, 11ßHSD1 is expressed in cardiomyocytes and in cardiac fibroblasts, as well as in neutrophils and macrophages recruited to the heart following MI (Gray, White, Castellan, McSweeney & Chapman, 2017). We have previously shown that 11ßHSD1 in cardiac fibroblasts plays a key role in the acute regulation of neutrophil recruitment after experimental MI in the mouse (Mylonas et al., 2017). However, while GCs are beneficial acutely in providing cardioprotection and suppressing inflammation, continued availability impairs wound healing, infarct repair (Fernandez-Ruiz, 2022; Huang, Yang, Chen, Liu, Chang & Chuang, 2013) and potentially cardiomyocyte regeneration (Sethi et al., 2023; Tao et al., 2020). Intervening with exogenous GCs in the post-MI period is detrimental to recovery (Huang, Yang, Chen, Liu, Chang & Chuang, 2013; Libby, Maroko, Bloor, Sobel & Braunwald, 1973). GCs regenerated by 11ßHSD1 are known to suppress angiogenesis (Small et al., 2005) and in a mouse model of MI suppression of *Hsd11b1* expression enhanced angiogenesis, promoted macrophage polarisation, reduced infarct expansion (McSweeney et al., 2010) and prevented the development of HF (White et al., 2016).

Drugs to inhibit 11ßHSD1 initially developed to treat diabetes (Anderson & Walker, 2013) and cognitive decline (Webster et al., 2017) have safely reached Phase 2 clinical trials. Although many 11ßHSD1 inhibitors (11ßHSD1i) have not thus far been developed further for clinical use, recent studies showing effective engagement with brain 11ßHSD1 in Alzheimer’s patients (Villemagne et al., 2024) and promotion of diabetic skin wound healing (Wilton-Waddell et al., 2024) continue to support their therapeutic promise.

The primary aim of the current study was to determine the potential for repurposing of 11ßHSD1i for use in the acute post-MI period to prevent the subsequent structural and functional remodelling that leads to HF. The study additionally investigated the impact of 11ßHSD1i intervention on neovascularisation, inflammation and extracellular matrix remodelling during wound repair. Pharmacological inhibition of 11ßHSD1 was initiated 48h after MI, by oral administration of the Astra Zeneca compound AZD8329 (Scott *et al*., 2012) in a translational mini-pig model of percutaneous balloon MI with reperfusion. To enhance the clinical relevance of the study all mini-pigs in both the vehicle and 11ßHSD1i treatment groups concurrently received drugs used therapeutically following post-MI: anti-platelet drugs (<u>aspirin</u>, <u>clopidogrel</u>), a statin (<u>simvastatin</u>) and an ACE inhibitor (<u>ramipril</u>).

## 2. METHODS

The *in vivo* animal regulated procedures were conducted in line with UK Home Office regulations (Animal Act 1986) at the MHRA compliant Translational Biomedical Research Centre - TBRC (Bristol, UK). The procedures conformed to the Directive 2010/63/EU and were undertaken under a Project License (7008975) granted by the Home Office after formal review and approval by the University of Bristol Animal Welfare and Ethics Review Body (AWERB). The reporting of data from these animal experiments is in keeping with the ARRIVE (Animals in Research: Reporting In Vivo Experiments) guidelines.

Adult female Goettingen mini-pigs (18-20kg) were supplied by Ellegaard, Denmark. Female mini-pigs were selected for this study as acute survival is improved relative to males following MI and reperfusion. On arrival at the University of Bristol, animals were housed for acclimatisation under regulated procedures in standards pens containing at least two individuals on a 12-h light/dark cycle with water and food provided *ad libitum*.

### 2.1 Drug treatment

All pigs received drug treatment prior to and after surgery as shown in Supplementary Figure 1, additional drug treatment used during surgery only is detailed in 2.2 below. Doses were selected based on previous studies in pig models. Animals were pre-treated with aspirin (75mg) and simvastatin (1mg/kg/day) in food (Accord, Barnstaple, UK) for 5 days prior to surgery and then daily until termination. Treatment with ramipril (2.5mg/day) and clopidogrel (75mg/day) in food was begun on the day after surgery and then daily until termination. On day 2 after MI, following MRI (as below), pigs were randomized according to a blind pre-defined sequence to receive either AZD8329 or vehicle daily until study termination. The drug vehicle was prepared by dissolving 11g of kleptulose powder (Roquette, France) in 110ml of working stock (90% apple juice and 10% water), pH=8, then mixed with 30g of oats to make a porridge. For drug treatment AZD8329 was dissolved at 50mg/kg of animal weight in the kleptulose solution at pH=10. Once the drug was dissolved, the pH was set to the original pH=8, prior to mixture with oats. Animals were trained daily to eat the porridge formulation without drug during the pre-surgery acclimatisation period. The formulation/ oats mixture was given to the animals on an empty stomach and entire feed consumption was monitored and recorded by the dedicated senior veterinary technician. The animals were not fed with their routine feed until 1 hour after the mixture had been consumed to ensure no interference with uptake of the drug. The dose of AZD8329 (50mg/kg/day) and formulation for delivery was selected following previous pharmacokinetic studies and modelling, with the aim of maintaining a plasma concentration at least 3 times the *in vitro* IC_50_ (0.03x10^-6^M) at steady state. The final plasma concentration of AZD8329 achieved, as well as steroid concentrations (Denham et al., 2024), were assessed in plasma collected at study termination by LC-MS/MS at the Edinburgh Clinical Research Mass Spectrometry Core Facility, as described below.

### 2.2 Percutaneous balloon MI and reperfusion

On the day of surgery mini-pigs received intramuscular premedication <u>ketamine</u> (10 mg/kg Vetoquinol UK Limited, Towcester, UK) and <u>dexmedetomidine</u> (15 µg/kg, Vetoquinol UK Limited, Towcester, UK). General anaesthesia was induced with <u>propofol</u> (incremental doses of 1 mg/kg, Zoetis UK Limited, London, UK) followed by intubation and ventilation, while anaesthesia was maintained with 1–2% <u>isoflurane</u>. Mechanical ventilation consisted of fraction of inspired oxygen, 0.21-1; tidal volume, 10 ml/kg; respiratory rate, 20 breaths per min; positive end-expiratory pressure, 0-3 cmH_2_O. Following IV heparinisation to keep activated coagulation time (ACT) >250sec, animals were subjected to closed chest MI procedure of the antero-apical LV territory under fluoroscopic guidance (Artis Zee, Siemens, Germany). Central vascular access was achieved with a two-way catheter sited in the left jugular vein and an introducer in the left carotid artery. Ringer’s lactate (4 ml/kg/h, B.Braun Melsungen AG, Melsungen, Germany) was used IV to maintain euvolemia. Continuous invasive blood pressure and routine 12 leads ECG were implemented throughout the MI procedure along with peripheral capillary oxygen saturation (SpO_2_) that was monitored by a pulse oximeter sited on the tail, with data capture integrated into the Datex-Aisys anaesthetic machine. Following instrumentation, the animals were allowed to stabilise for a minimum of 30 minutes. Animals were then taken to the adjacent interventional theatre. To minimise the risk of ventricular fibrillation (VF), during/after the MI procedure infusions of <u>amiodarone</u> (150 mg) and magnesium sulphate (4 mmol) were administered over 90min via the jugular venous line, commencing soon after access was gained.

Following left coronary angiography, a 0.014-inch interventional guide wire (Runthrough floppy, Terumo, Tokyo, Japan) was steered to the distal left anterior descending (LAD) coronary artery, and an appropriately sized balloon (2.5-3.0 mm Emerge, Boston Scientific, Marlborough, MA, USA) was positioned beyond the LAD first diagonal branch. The balloon was then inflated to stop blood flow to induce ischemia and then kept up for 60min to induce acute MI. Blood pressure was kept within reference intervals using small increment of metaraminol (0.5mg) IV and/or volume replacement while SpO_2_ levels were kept >94%. ON completion of 60min of ischemia, the balloon was deflated to start reperfusion and animals closely monitored for a period of 45-60min till fully awake and deemed suitable for transfer to the maintenance area according to predefined standard operating procedure by senior expert veterinary anaesthetists. Amiodarone infusion (150mg over 90min) was started before coronary occlusion to prevent ventricular fibrillation (VF) with additional bolus (100mg) and direct current (DC) cardioversion used in case of VF occurrence during ischemia. In case of VF or haemodynamic compromise during the ischaemic period, resuscitation through direct current (DC) cardioversion, manual chest compression and/or inotropic support was provided, as required according to a predefined protocol. If pre-defined resuscitation cycles were unsuccessful, the experiment was terminated. Surviving animals were recovered/monitored for 45-60min before being returned to their pen.

### 2.3 Cardiac MRI

On day 2 after MI baseline characterisation of LV function, volumes and MI scar size was collected under general anaesthesia via magnetic resonance imaging (MRI). Briefly, animals surviving the MI procedures were re-anaesthetized 2 days later and received intravenous <u>pancuronium bromide</u> (0.2 mg kg^-1^ bolus followed by 0.1 mg kg^-1^ h^-1^) to enable breath holds during cardiac MRI image acquisition, achieved by temporary interruption of ventilation. Images were acquired on a Magnetom Prisma® 3T system, (Siemens Healthcare limited, Erlangen, Germany). Cardiac MRI scans were performed in supine recumbence with the surface coil positioned centrally on the sternum. Images were acquired on a Magnetom Prisma® 3T system, (Siemens Healthcare limited, Erlangen, Germany) using the integrated spine coil posteriorly, and the Siemens ‘Body 13’ coil anteriorly. Cardiac gating was achieved using a surface ECG or, in cases where there was poor triggering, a peripheral pulse monitor placed on the tail. Due to the importance of a stable ECG, the animals were shaved, and the skin prepared with Nuprep® (Weaver and Company, Aurora, CO, USA) to improve skin contact. After acquisition of localiser images, True–FISP functional cinematic images were acquired in the three long axis planes (2, 3- and 4-chamber orientation) of the left ventricle, and in a contiguous stack of short axis slices from base (at the level of the mitral valve) to the apex. The acquisition of parameters was in keeping the established protocols allowing visual assessment of the 17 American Heart Association (AHA) myocardial segments. After cine imaging, a dose of 0.2 mmol kg^-1^ body weight of Gadobutrol (Gadovist^TM^, Bayer, Newbury, Berks, UK) contrast agent was administered, late gadolinium enhancement (LGE) images were acquired after 8-10 minutes in line with established guidelines (Kramer, Barkhausen, Flamm, Kim, Nagel & Society for Cardiovascular Magnetic Resonance Board of Trustees Task Force on Standardized, 2013) to measure scar size. Images were acquired in the long axis and short axis using a 2D FLASH PSIR pulse sequence: slice thickness, 8 mm; base resolution, 256; phase resolution, 64%; TE, 1.55 ms; TR, 750 ms; TI, 300-350 ms. On completion, all animals were recovered for 45-60min and then sent back to their maintenance pen. Imaging was performed by a blinded expert radiographer (CJ). On completion of the 4-week study period, animals underwent a repeat cardiac MRI as above, followed by controlled termination, as below. Subsequent blinded analysis of cine and LGE images was performed at the Edinburgh Clinical Research Imaging Centre using commercial clinical analysis software cvi42 (Circle Cardiovascular Imaging).

### 2.4 Blood sampling, troponin and LC-MS/MS for steroids and drug concentrations

Blood was collected under GA or during recovery from jugular or peripheral vein line at baseline, end of ischemia and at 1min, 1h, 24h, 48 and 28d post-MI. 10 mL of blood was collected into EDTA vacutainers, then centrifuged at 1000 g for 10 min at 4°C, plasma was collected and stored at -80°C. Serum was collected at baseline, 24h, 48h and 28d post-MI by inverting 10 mL of blood 5 times, leaving at RT for 39 min to allow clot formation and removing clot by 1500g centrifugation for 10 minutes at 4°C before storing serum at -80°C. Only 7 serum samples from each group were suitable for troponin analysis due to evidence of haemolysis in some samples. All subsequent analysis was carried out in a blinded manner. Troponin I was assessed at the TBRC in plasma samples by commercial ELISA assay according to the manufacturer’s instructions (Life Diagnostics, UK). The steroids <u>cortisol</u> and <u>cortisone</u> were measured at the Edinburgh Clinical Research Mass Spectrometry Core Facility in pig plasma as described in detail by Denham et al (2024). In short plasma (200 µL) was extracted using supported liquid extraction and analysed by targeted LC-MS/MS. The glucocorticoid (cortisol and cortisone) concentrations were calculated using linear regression of the calibration curve, as described previously (Denham et al., 2024). To measure the circulating concentration of the drug AZD8329, plasma samples were pre-diluted 1000 x in water (LC-MS grade, VWR, 83645.320) and 100 µL of these diluted samples were extracted alongside calibration standards covering range 2.5–1000 ng/mL of AZD8329, spiked into 100 µL of water (LC-MS grade). Samples and calibration standards including blanks, were pipetted into 2 mL deep 96-well collection plate (Biotage, 121-5203), sealed using 96-well plate sealing film (VWR, 391-1250) and shaken for 10 min at 600 rpm on a 3005 Plate Shaker (GFL) to allow mixing. The collection plate and PPT+ plate were both loaded into an Extrahera liquid handling robot (Biotage, Sweden), which loaded 400 µL acetonitrile + 0.1% v/v formic acid (VWR, 83640.320 and Fisher, 10596814, respectively) followed by transfer of the samples (100 µL) into the PPT+ plate. After a 10 min incubation at RT, positive pressure was applied to the PPT+ plate to elute samples into a 96-well collection plate. The collection plate was transferred to a TurboVap 96 Dual Evaporator (Biotage, Sweden) with an oxygen free nitrogen gas flow of 30 L/min and the samples reduced to dryness at 40°C. Once dry, the samples and standards were dissolved in 10% v/v acetonitrile (100 µL). The plate was sealed and shaken for 10 min at 600 rpm to ensure all analytes resolubilised. The reconstituted samples were subject to LC-MS/MS analysis, beginning with separation on a Waters Acquity I-Class ultra-performance liquid chromatography system (Waters, UK). The separation was carried out on a Kinetex C18 column (50 x 2.1 mm; 1.7 µm), (00B-4475-AN; Phenomenex, UK), maintained at 50°C with a flow rate of 0.4 mL/min and a mobile phase A (MPA) – water with 0.1% v/v formic acid; mobile phase B (MPB)– acetonitrile with 0.1% v/v formic acid. Starting at 5% MPB for 1 min, increasing to 100% MPB at 3.5 min, held for 2 min before re-equilibrating to 5% MPB over 2 mins, giving total run time of 7.5 mins per sample. Detection of analyte (11βHSD1i, AZD8329) was performed using positive ion electrospray ionisation mode on a QTRAP 6500+ mass spectrometer (Sciex, UK) operated by Analyst 1.7.1. The source was operated at 600°C with an IonSpray voltage of -4.5 kVcurtain gas of 40 psi, nitrogen nebuliser ion source gas 1 (GS1) of 60 psi, and heater ion source gas 2 (GS2) of 40 psi. AZD8329 eluted at 3.6 mins and was monitored by using the multiple reaction monitoring transition; *m/z* 422.07 è 135.1 (declustering potential 216V, collision energy 39V and collision exit potential 16V). Quantification of AZD8329 was performed using linear regression of the calibration curve using MultiQuant v3.0.3 (Sciex, UK) for calculation of the concentration of AZD8329 in plasma.

### 2.5 Controlled termination and tissue collection

On completion of follow-up cardiac MRI acquisition, animals were taken to the TBRC interventional room for a controlled termination. Under general anaesthesia and continuous monitoring of vital parameters (BP, ECG, SpO_2_ and temperature, a median sternotomy was performed, and <u>heparin</u> administered to achieve an activated coagulation time >250 s. Next, the superior and inferior vena cava and the aorta were clamped in sequence and 500 ml of cold (4 °C) cardioplegia solution was administered into the aortic root at high pressure (>200 mmHg) to achieve diastolic standstill. This solution included magnesium chloride (1.193 g), potassium chloride (1.193 g), and <u>procaine</u> hydrochloride (272.8 mg) per 20 ml at clinical grade, diluted into 1L of normal saline (Martindale Pharmaceuticals, Romford, Essex, UK). At termination, the whole arrested hearts were rapidly harvested and submerged in cold cardioplegia for tissue sampling as previously reported (Ascione et al., 2024). Three consecutive circular transverse sections of the left ventricle were dissected; the LV apical third section was flash-frozen as a whole at maintained at -80°C for later cryo-sectioning and MSI; the LV mid third section was promptly divided in undamaged, transition zone, peri-infarct zone, and scarred zone, each aliquoted, flash-frozen and kept at -80°C for transcriptomics and proteomics. The LV basal third section was fixed whole in formalin for 24 hours, and transferred in 70% ethanol and kept at 4°C. All heart specimens were packed at clinical grade and couriered to the University of Edinburgh further evaluation a below.

### 2.6. Immunohistochemistry

Cross-sections through the whole left ventricle that included the infarct, border zone and non-infarcted myocardium were dissected, fixed in 10% formalin for 24h and subsequently maintained in 70% alcohol for further processing and immunohistochemical staining of CD31, α-smooth muscle actin (αSMA) CD163, fibroblast activation protein (FAP) and wheat-germ agglutinin (WGA). All samples were blinded prior to analysis. A number of samples had to be excluded from analysis because of incomplete fixation, this resulted in a reduced number in both groups relative to the *in vivo* data. The sections were processed to wax blocks, sectioned at 4 µm thickness and mounted on Epredia™^T^ SuperFrost Plus™ adhesion slides (Fisher Scientific, 10149870).

ImmPRESS® HRP Horse Anti-rabbit (MP-77452) or the Goat Anti-mouse (MP-7401) IgG Polymer kits from Vector Laboratories was used for all IHC staining. After de-paraffinisation and rehydration, antigen retrieval was carried out by submerging slides in 10 mM citric acid (pH 6.0, Sigma) and incubating in a pressure cooker for 15 min. Slides were then washed in water, dried and incubated with endogenous peroxidase blocking solution Bloxall® (Vector Laboratories, SP-6000-100) for 10 min at RT. After washing in PBS, sections were blocked with 2.5% horse or goat serum for 20 mins at RT. The slides were then incubated with rabbit anti-CD31 (Abcam, ab28364, 1:500 in PBS), both anti-CD31 and mouse anti-αSMA (Sigma, A2547, 1:5,000 in PBS), rabbit anti-FAP (Abcam, ab207178, 1:500 in PBS) or mouse anti-CD163 (Novus Biological, NB110-40686, 1:500 in PBS) overnight at 4°C. Slides were washed with PBS for 5 min, followed by incubation with HRP polymer-conjugated anti-rabbit or anti-mouse secondary antibody for 30 min at RT. Slides were washed with PBS twice for 5 min and incubated with DAB stain (ImmPACT® DAB Substrate, Peroxidase, SK-4105) for 2-5 minutes, until colour uptake was observed. After counterstaining with haematoxylin, slides were dehydrated and mounted with xylene-based mounting media. Images were generated by scanning slides on the AxioScan scanner (ZIESS), and quantification was performed using QuPath (Bankhead et al., 2017). To measure vessel density, vessels were counted in 10 random 1.6 mm^2^ boxes generated in the infarct area, border zone and remote myocardium, and categorised based on their diameter (<10 µm: capillaries, 10-100 µm: medium-sized arteries, >100 µm: large arteries). For FAP, area was calculated based on a threshold of DAB intensity in 20 random 1.6 mm^2^ boxes generated in the infarct area. For CD163, QuPath’s cell detection module was applied on 10 random 1.6 mm^2^ boxes in the infarct area.

To measure cardiomyocyte cross-sectional area (CSA), cell membranes in heart sections were stained with WGA. After antigen retrieval, slides were incubated with 2.5% horse serum for 20 min at RT. Slides were then incubated with Isolectin B4 (1:100, biotinylated, Vector) for 1h at room temperature to stain endothelial cells. After washing in PBS for 5 min, slides were incubated with secondary detection reagent for IB4 (1:200 streptavidin-Alexa Fluor 555, Sigma, S-32355) and WGA-Alexa Fluor 488 (1:200, ThermoFisher, W11261) for 1h at RT. Slides were washed with PBS and counterstained with DAPI (1:500) to stain nuclei for 10 min at RT. After a quick wash in water, slides were mounted using PermaFlour mounting medium (FisherScientific, 12695925). Images were taken on the Axiovert 200M of at least 8 fields of the remote myocardium in each section. Fields were confirmed to be on the transverse plane by presence of vessels stained with the IB4. Cardiomyocyte cross-sectional area of at least 10 cardiomyocytes/field was measured using QuPath and averaged over all fields assessed.

### 2.6 Picrosirius red histological staining and polarised light analysis

Fixed LV cross-sections (4 µm thickness) were stained with picrosirius red (PSR) for collagen and polarised light analysis. After de-paraffinisation and rehydration, sections were incubated in 0.5% picrosirius red solution (Sigma-Aldrich) for 2 h at RT. Collagen area was quantified in the infarct using the QuPath threshold function in 20 random 1.6 mm^2^ boxes. To analyse collagen organisation, sections were imaged with polarised light using an Axiovert 200M microscope (Carl Zeiss Ltd) and 3-5 random fields in the infarct were imaged at magnification of 40x. Collagen has natural birefringence that is enhanced by PSR, causing its fibres to appear either red or green under polarised light. Thin, newly synthesised fibres tend to appear yellow-green (YG), while thick, established fibres appear red-orange (RO). The areas of YG and of RO collagen were determined using the Threshold Colour module in ImageJ. Based on optimisation in trial images, the hue threshold for RO was set between 0-35, while the hue threshold for YG was set between 36-110 (RGB hue). Brightness was consistently set at a range of 15-255 and saturation at 0-255. Using the threshold to detect the appropriate colour hue, the image was then filtered for the selected area and threshold colour transformed to black and white. The area of the detected colour was then generated in pixels and calculated in µm^2^ based on image pixel size.

### 2.7 MALDI Mass Spectrometry Imaging to localise AZD8329 in the myocardium

A LV cross-section that included the infarct, border zone and non-infarcted myocardium was dissected from each heart at study termination and flash frozen by floating on liquid nitrogen atop an aluminium foil boat. Cross-sections were blinded for identification and sectioned into 10 µm thick slices using a cryostat (CryoStar NX50, Thermo Scientific) at -18°C and thaw mounted onto indium tin oxide (ITO)-coated glass slides (Delta technologies, CG-81IN-S115). Before applying the MALDI matrix, slides were left to dry in a vacuum desiccator for 10 min. Matrix and solvent were prepared fresh before every experiment by dissolving of 1,5-diaminonapthalene (1,5-DAN, 5 mg/mL) in 50% v/v acetonitrile 10 coats of the matrix were applied in a crisscross pattern by the HTX M3 TM-Sprayer™ (HTX Technologies) at 0.1 mL/min flow rate with 2 s dry time between coats. The nozzle temperature was set at 75°C and the spraying was conducted at 12 mm/min velocity with a gas flow rate of 2L/min.

Mass spectrometry imaging was performed using a 12T SolariX MALDI-FTICRMS (Bruker), employing a Smartbeam laser (1 kHz). The instrument was operated with SolariX control v2.2.0 (Build 150) and fleximaging v5.0 (Build 89). Analysis was performed in negative ion mode, using 1000 laser shots with a width of 150 µm, frequency of 2000 Hz and power of 35%. Ions were isolated using continuous accumulation of selected ions (CASI) mode Q1 mass of 420 mass-to-charge ration (*m/z)* and an isolation window 100 *m/z*. Analysis was performed using Fleximaging v5.0 software (Build 89). Data were loaded and processed at 8000 data points and ion cyclotron resonance noise reduction threshold of 95%. A mass defect limit of 0.025 Da and minimum intensity of 2% were used to filter spectra. Images were produced by selecting 11βHSD1i peak in the spectra at 420.2265 ± 0.002 Da. The same slide was washed of matrix and stained with H&E to allow identification of infarct.

### 2.8 Proteomics

9 frozen LV tissue blocks were included from each treatment group, this represented all of the vehicle group and 9/11 of the 11βHSD1i treatment group. The 2 samples that were excluded were from the pigs with the least improvement in LV function with 11βHSD1i treatment. LV tissue was blinded for identification and homogenised prior to protein precipitation for proteomic analysis using Ultra-High-Performance Liquid Chromatography/Mass spectrometry (UHPLC/MS). Tissue blocks were transferred to 2 mL tubes pre-filled with 2.8 mm ceramic beads (Fisher Scientific, 15565799) and homogenised twice in 1 mL PBS (without Ca^2+^ or Mg^2+^), containing 1% protease inhibitor cocktail (v/v, Sigma-Aldrich, P8340) on the Precellys 24 tissue homogeniser (Bertin Instruments) at 6500 rpm for 50 s. Samples were allowed to cool on ice for 5 min between cycles. Homogenised samples were then centrifuged at 14000 g for 10 min at 4°C. The supernatant was discarded, and the remaining protein pellet was washed in PBS and centrifuged at 14000 g for 10 min at 4°C. Protein pellets were resuspended in lysis buffer (6 M Guanidine hydrochloride, 100 mM Tris pH 8.5 10 mM TCEP 15 mM chloroacetamide) with probe sonication. Lysates were then heated to 95°C for 5 min, followed by a digestion with 1 µg LysC (Wako, Japan) for 4 h at 37°C. After diluting 6x with LC-MS-grade water, 1 µg of porcine trypsin (Pierce, UK) was added and samples were digested over night at 37°C. Samples were acidified using 0.5% trifluoracetic acid and clarified by centrifugation at 18000 × g for 15min. They were then desalted using C18 stop-and-go extraction (Stage) tips made in house (adapted from (Rappsilber, Ishihama & Mann, 2003)). Two disks were cut from C18 solid-phase extraction material (supplier) using an 18-gauge blunt-ended needle. The disks were then placed on top of each other inside a 200 μl EasyLoad pipette tip (Greiner Bio-One), which were loaded into a custom-built tip holder over a deep 96-well plate. C18 Stage tips were washed with methanol by centrifugation at 300 g for 2 min, and then equilibrated with 0.1% TFA and centrifuged at 500 g for 5min. The acidified peptides were then eluted through the C18 Stage tips with 80% ACN containing 0.1% TFA and centrifuged at 200 g for 5 min. Peptide concentration was measured using Nanodrop. The de-salted peptides (2 µg) were loaded into a 50 cm emitter packed with 1.9 µm ReproSil-Pur 200 C18-AQ (Dr. Maisch). Peptides were separated by a 140 min linear gradient from 5-30% ACN containing 0.05% acetic acid on an UltiMate 3000 UHPLC Nano (Thermo Fisher Scientific) coupled to Orbitrap Fusion Lumos Tribrid mass spectrometer (Thermo Fisher Scientific). The mass spectrometer was operated in DIA mode, acquiring *m/z* between 35-1650 Da at mass resolution of 120,000. The peptides acquired were then further fragmented by MS/MS on 45 *m/z* windows between 200-2000 Da with 0.5 Da overlap at normalised collision energy of 27 and a resolution of 30,000. The generated RAW spectra files were quantified using DIA-NN (Demichev, Messner, Vernardis, Lilley & Ralser, 2020). The data were searched against the UniProtKB Sus scrofa database (Taxonomy ID 9823), from which an in-silico spectral library was generated via the software. The database and spectra RAW files were searched for peptides 7-30 residues in length with a maximum of one missed trypsin cleavage and including modifications introduced during sample preparation (*N*-terminal methionine excision and cysteine carbamidomethylation). Precursor FDR was set at 1% and the quantification strategy at “Robust LC” for high precision. In each search, the neural network classifier was run in double pass mode to reduce interference. DIA-NN generated a list of proteins and their relative intensities that was used for further differential analysis using the *limma* R/Bioconductor package (Ritchie et al., 2015). The list of differentially expressed proteins generated from *limma* analysis is in Supplementary Table 6.

To identify enriched pathways and their underlying protein-protein interactions, the list of differentially regulated proteins was analysed further using Reactome (Fabregat et al., 2017) and STRING (Szklarczyk et al., 2019). Pathways and interactions were projected to the human proteome. Enriched pathways were identified by mapping differentially expressed proteins to *Reactome* pathways which used a hypergeometric test to assess over-representation. To control for multiple testing, Benjamini-Hochberg correction was applied and pathways with a corrected p-value below 0.05 were considered significantly enriched. The list of significantly enriched pathways generated from Reactome pathway analysis tool is provided in the Supplementary Table 5. STRING predicted protein-protein interaction in the differentially expressed proteins using a database based on various evidence sources, including co-expression, gene fusion, text mining, and experimental data. The fold change data generated using *limma* were superimposed on the network generated. STRING operation and visualisation was conducted on Cytoscape v3.10.0 (Shannon et al., 2003).

### 2.9 Real-time polymerase chain reaction

Flash frozen LV tissue blocks from the infarct and border zones were blinded for identification and homogenised using a 5 mm stainless steel bead (Qiagen, 69989) and TissueLyserII (Qiagen) at 30 Hz for 1.5 min 2-5 times. RNA was extracted using the RNeasy Fibrous Tissue Mini Kit (Qiagen, 74704) following the manufacturer’s protocol. RNA was quantified using a Nanodrop. Based on average of 2 measurements, the volume required to produce 1 μg of cDNA was calculated. Reverse transcription was conducted using the High Capacity RNA-to-cDNA kit (ThermoFisher, 4387406) following the manufacturer’s instructions. All samples were pipetted into MicroAmp Optical 96-well plate (Applied Biosystems, N8010560) and incubated in a Veriti 96 Well Thermal Cycler (Applied Biosystems) at 37°C for 1 h with a cover temperature of 105°C. The reaction was stopped by heating the plate to 95°C for 5 min. To perform qPCR, FAM-labelled TaqMan Gene Expression assays (ThermoFisher, Supplemental Table 1) were used. cDNA was diluted to approximately 33 ng per µLof nuclease-free water (Qiagen, 129114). The TaqMan assay and TaqMan Fast Advance Master Mix (ThermoFisher, 4444556) were prepared as described in Supplemental Table 2 and pipetted in triplicates in a 384 well lightcycler plate (Starstedt, 72.1985.202). The plate was then sealed using adhesive sealing sheets (ThermoScientific, AB-0558), flipped to mix, and centrifuged at 4000 rpm for 1 min at 4°C and run on the LightCycler 480 II (Roche) using the steps outlined in Supplemental Table 3.

### 2.10 Data and Statistical Analysis

Power analysis prior to the study (based on previous data from Bristol in adult farm pigs) identified a requirement of n=11 in each group to provide 90% power to detect a 5% change in left ventricular ejection fraction, with a significance level of P<0.05. All analysis was completed in a blinded manner. Group sizes in some figures were reduced where it was not possible to collect *in vivo* images (for LGE MRI), where blood samples were haemolysed, or where LV tissue had been incompletely fixed. Only selected samples were included in mass spectrometry and proteomic analysis due to assay costs, selection was on the basis of representative LVEF in the *in vivo* study. Statistical analysis was performed using GraphPad Prism 9.2.0. Two-way ANOVA with Bonferroni post-hoc testing was used when 2 variables were being compared, while an unpaired t-test was used to analyse single variables being compared across the two treatment groups. All data are presented as mean±SEM, and significance set at P<0.05. Statistical analysis for differential expression analysis on proteomics was conducted using the *limma* R/Bioconductor package (Ritchie et al., 2015).

## 3. RESULTS

### 3.1 Surgical outcomes

In total 25 animals were entered into the study and randomly allocated to the two treatment arms (*Supplementary* Figure 1). 4 pigs died due to intractable VF/VT during the MI procedure. A further pig was excluded due to lack of infarct scar on initial MRI imaging, and no loss of function suggesting that coronary occlusion was unsuccessful. This resulted in 9 pigs in the vehicle treatment group and 11 the 11ßHSD1i treatment group.

### 3.2 The 11ßHSD1 target is successfully engaged by AZD8329

LC-MS/MS analysis of blood collected at study termination (24h after the final oral drug treatment) revealed a plasma AZD8329 concentration of 92±14µM in treated pigs. The drug was not detected in vehicle-treated pigs. The plasma level of the 11ßHSD1 substrate cortisone was significantly elevated in treated pigs, 23.8±2.4nM compared to vehicle treated animals, 16.3±1.2nM (P<0.05), consistent with inhibition of enzyme activity.

### 3.3 Administration of an 11ßHSD1i after MI prevents functional decline and adverse ventricular dilatation

Late Gd enhancement MRI imaging (*Figure 1a*) shows that infarct size was equivalent in both treatment groups prior to initiation of treatment with 11ßHSD1i or vehicle (*Figure 1b*). This was confirmed by equivalent concentrations of troponin I in blood collected after induction of MI and during recovery (*Figure 1c*). Cardiac function, assessed by LVEF and cardiac output, was also equivalent in both treatment groups at 48h after MI and prior to initiation of treatment with 11ßHSD1i (*Figure 2a, Supplementary Table 4*). By 28 days later LVEF had decreased significantly in the vehicle treated group (P<0.05, *Figure 2a, b*) compared to the immediate post-MI period, but was unchanged in the pigs treated with 11ßHSD1i from 48h after MI (P<0.05). LVEF was increased by 9.6 ± 4.2% following 11ßHSD1i treatment. LVESV increased in vehicle treated animals compared to those receiving 11ßHSD1i (P<0.05, *Figure 3b*) and the same pattern was seen for LVEDV (*Figure 3a*), although this failed to reach statistical significance (P=0.07). There was no influence of 11ßHSD1i treatment on myocardial mass or cardiomyocyte cross-sectional area (*Figure 3, c-f*).

**Figure 1.**
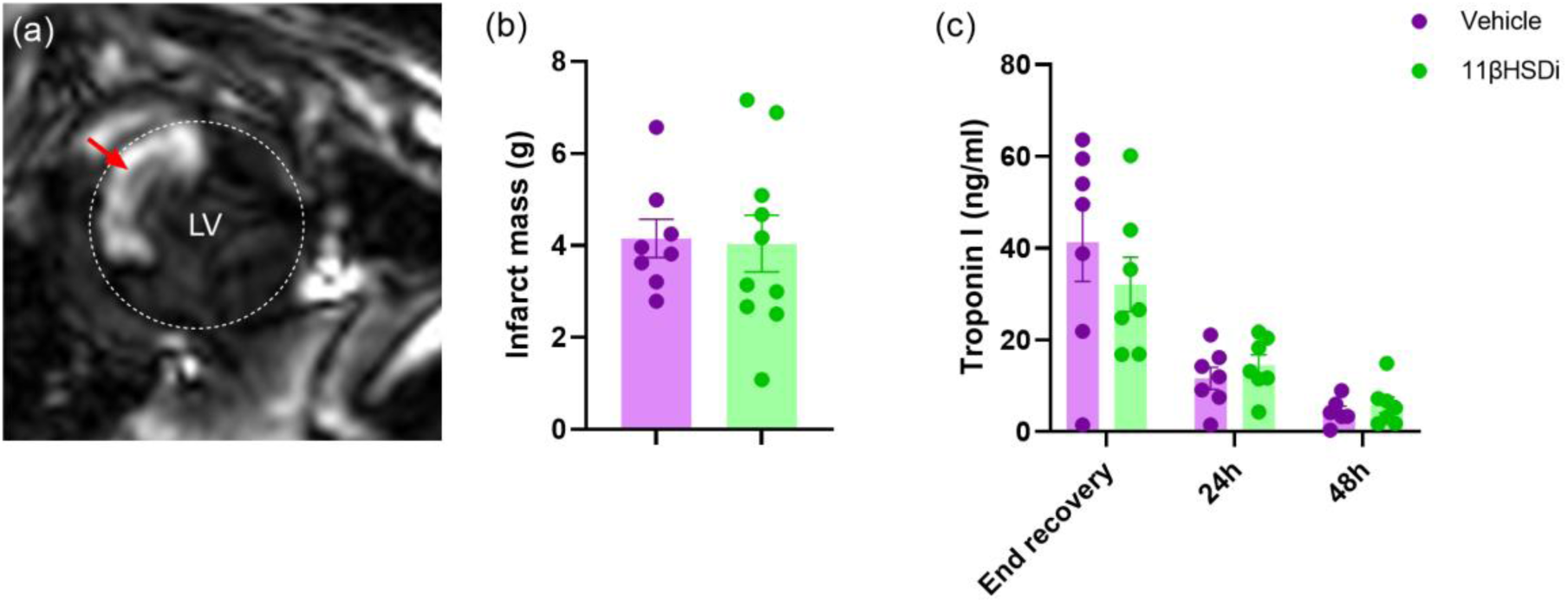
Infarct injury was equivalent in both study groups prior to initiation of treatment with 11βHSD1i or vehicle. Infarct injury was detected by late gadolinium enhancement magnetic resonance imaging (a, highlighted by red arrow) at 48h after MI (b), vehicle, n=8, 11βHSD1i, n=10, and confirmed by troponin I assay in blood samples collected immediately after surgery (end recovery) and at 24 h and 48h after surgery (c, vehicle n=7, 11βHSD1i n=7). Data presented as mean ± SEM, analysis was by two-way ANOVA.

**Figure 2.**
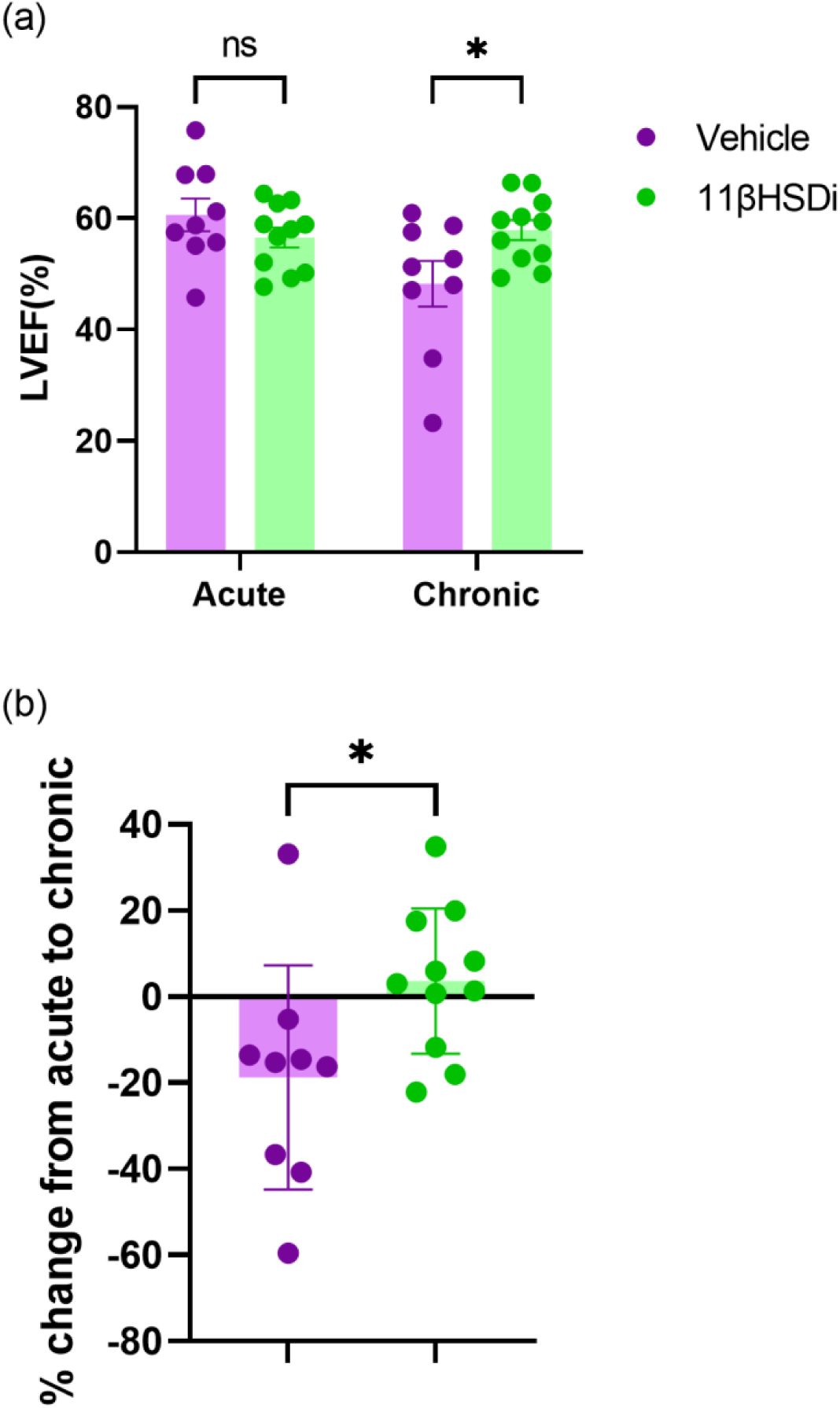
11βHSD1i therapy preserves cardiac function after MI, relative to vehicle. Left ventricle ejection fraction (LVEF), assessed by MRI in vehicle (n=9) and 11βHSD1i (n=11) treated pigs at 48h (Acute) and 28d (Chronic) after MI (a) and (b) the change between the 2 timepoints. Data are presented as mean ± SEM. *P<0.05, ns: not significant, two-way ANOVA with Bonferroni’s post-hoc test (a) or one-tailed unpaired t-test (b).

**Figure 3.**
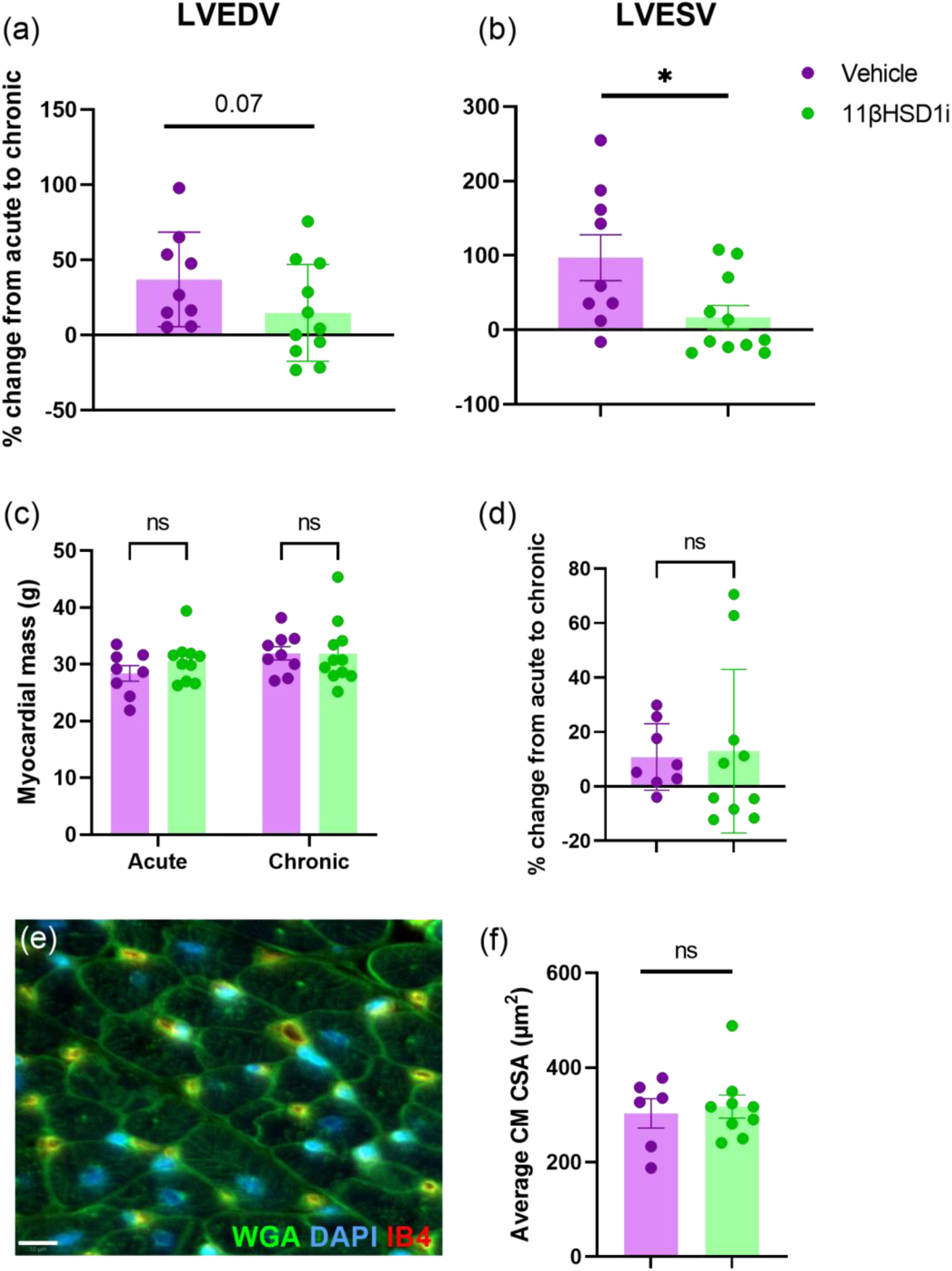
Intervention with 11βHSD1i therapy up to 28days post-MI reduces ventricular dilation but has no impact on cardiac mass or cardiomyocyte cross-sectional area. The difference in left ventricular end diastolic volume (LVEDV, a), end-systolic volume (LVESV, c) and myocardial mass (c) assessed by MRI between 48h and 28days after induction of MI in pigs treated with vehicle (n=9) or 11βHSD1i (n=11). Cardiomyocyte cross-sectional area (CM CSA e, f) assessed in immunostained ventricle collected 28 days after MI in both treatment groups (average of 130 CMs for each). Data are presented as mean ± SEM. *P<0.05, ns: not significant, two-way ANOVA with Bonferroni’s post-hoc (c) or one-tailed unpaired t-test (a, b, d, f).

### 3.4 Neither infarct size nor neovascularisation are influenced by 11ßHSD1i intervention

Neither LGE assessed infarct mass (*Figure 4 a,b*), nor histologically-derived infarct area (*Figure 4 c,d*) were influenced by 11ßHSD1i treatment after MI. The number of CD31^+ve^ capillaries was reduced in the infarct and border zones relative to the remote area of the left ventricle but 11ßHSD1i treatment had no effect on capillary density (*Figure 4 e,f*), or on the density of CD31^+ve^ αSMA^+ve^ smooth muscle coated vessels (*Figure 4 g,h*).

**Figure 4.**
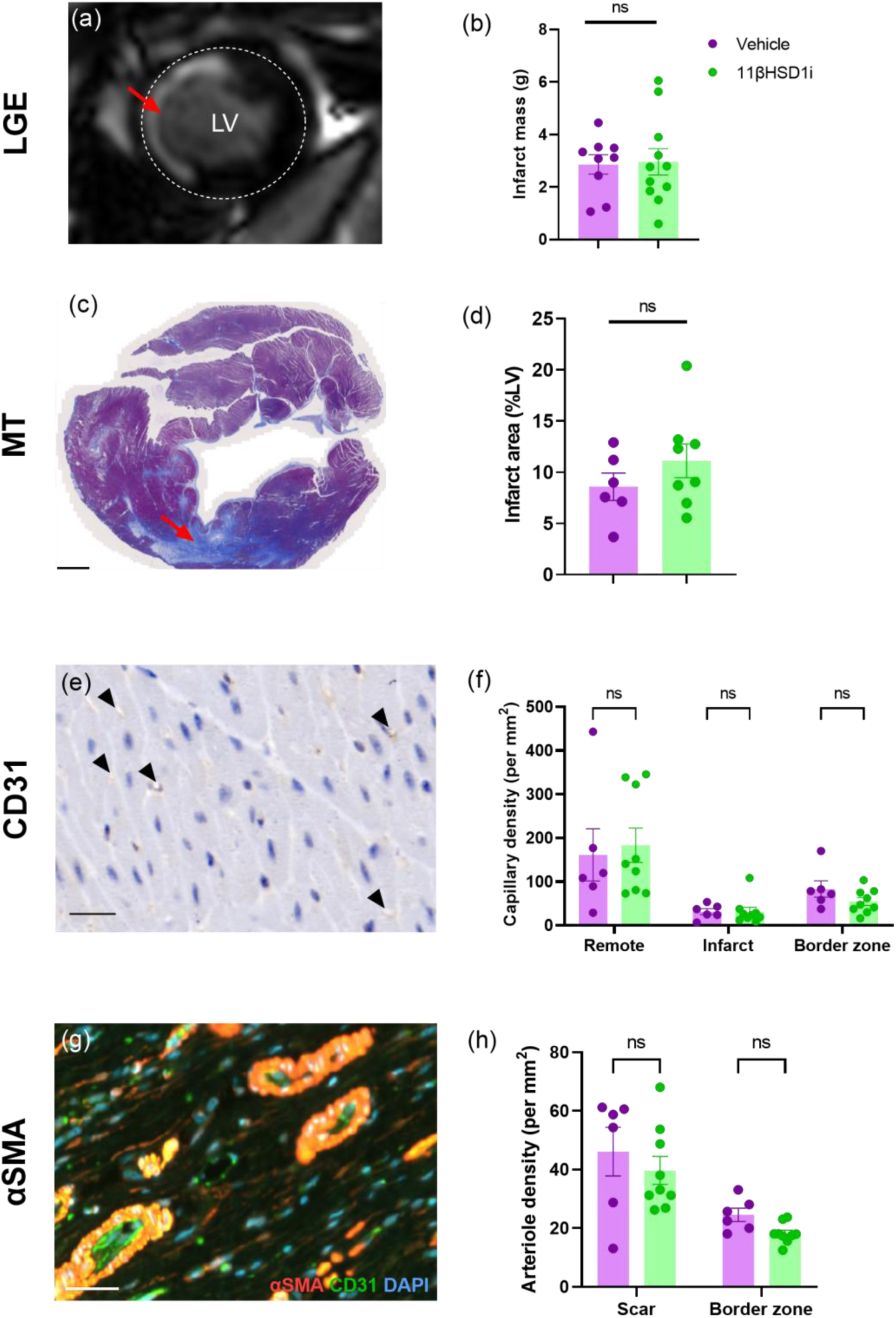
Neither infarct size, nor neovascularisation are impacted by 11βHSD1i therapy up to 28days post-MI. Infarct size measured using late-gadolinium enhancement MRI (a, b) and Masson’s trichrome (MT) histological stain (c, d) at 28d post-MI in pigs treated with vehicle (n=9 for MRI infarct mass and n=6 for histological infarct area) or 11βHSD1i (n=11for MRI infarct mass, and n=9 for histological infarct area). Vessel density of capillaries (CD31^+ve^, diameter <10µm, e) measured in remote, infarct and border zone (f) and arterioles (CD31^+ve^αSMA^+ve^, diameter 10-100µm, g) measured in the infarct and border zone (h). Red arrow: infarct, black arrows: capillaries. Scale bar: 2mm (c), 20µm (e) and 50µm (g). Data are presented as mean ± SEM. ns: not significant, two-tailed unpaired t-test (b, d) or two-way ANOVA (f, h).

### 3.5 11ßHSD1i AZD8329 is located in the repairing infarct scar after *in vivo* dosing and modifies collagen birefringence

Immunohistochemistry identified CD163^+ve^ macrophages (*Figure 5a*) and fibroblasts staining positive for fibroblast activation protein (FAP, *Figure 5b*) in the infarct zone relative to the remote myocardium, indicative of ongoing repair in the infarct zone 28 days after MI. The 11ßHSD1i drug AZD8329 administered *in vivo* until 24h prior to study termination was subsequently detected by MALDI MSI (*Figure 5c*) co-located with FAP (*Figure 5d*) in the scar, suggestive of intervention in the scar repair process. There was no influence of drug treatment on collagen content (*Figure 6a,b*) or on expression of COL1A1 mRNA or of COL3A1 mRNA (*Figure 6c*). The extent of CD163^+ve^ macrophage infiltration was also unaffected by 11ßHSD1i intervention (*Supplementary* Figure 2). Examination of picrosirius red staining under polarised light revealed that 11ßHSD1i treatment resulted in a relative increase in representation of narrow green-yellow collagen fibres compared to thicker red-orange fibres (*Figure 6d-f,* P<0.01). This was accompanied by a reduction in the expression of mRNA encoding the collagen cross-linking enzyme, lysyl oxidase (LOX, *Figure 6g*, P<0.05).

**Figure 5.**
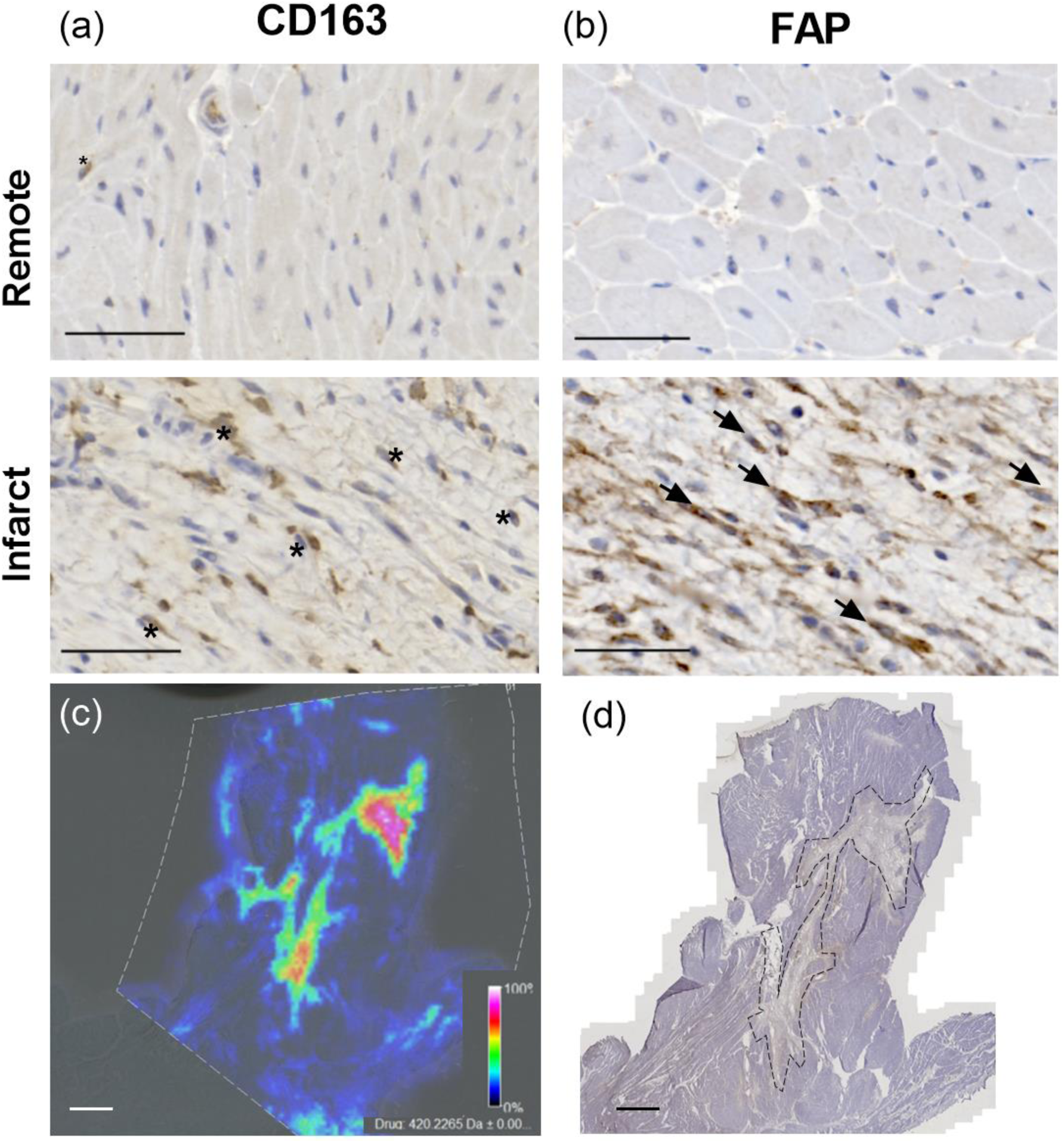
11βHSD1i is located in the repairing infarct scar after *in vivo* dosing. Representative images from CD163^+ve^ (a, stars) and fibroblast activation protein (FAP)^+ve^ (b, arrows) cells in the remote undamaged myocardium and infarct of vehicle (n=6) and 11βHSD1i (n=8) 28d post-MI. 11βHSD1i detection via MALDI imaging (c) corresponds with activated fibroblasts in the infarct area (d). Data are presented as mean±SEM. One-tailed unpaired t-test. Scale bar: 50µm (a-b), 2mm (c-d). Heat map (0% black to 100% white) represents intensity of 11βHSD1i signal at 420.2265±0.002 Da.

**Figure 6.**
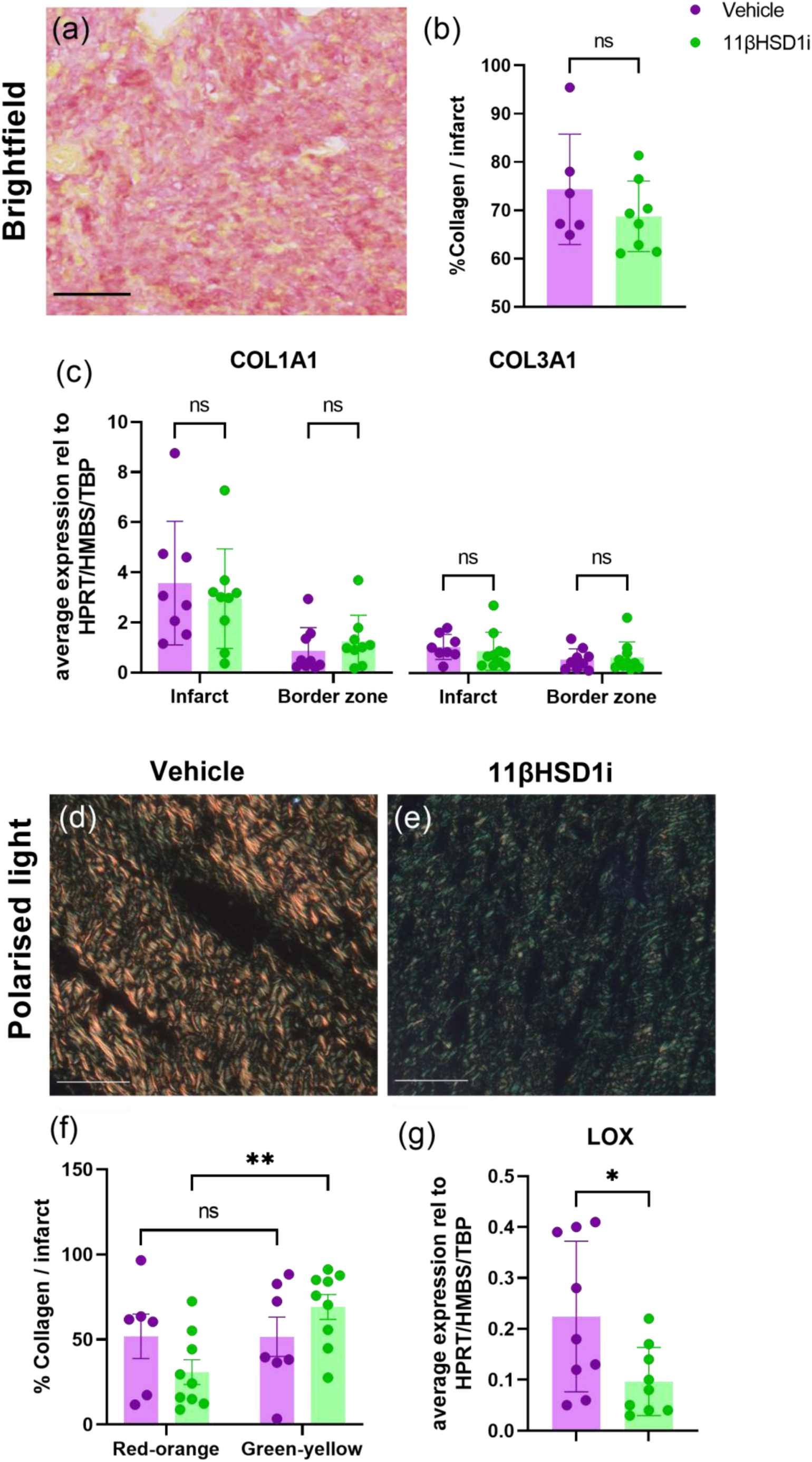
11βHSD1i therapy has no impact on collagen expression or content, but regulates organisation. Percent collagen in the infarct assessed by picrosirius red stain (a, b) and relative expression of COL1A1 and COL3A1 (c) in the infarct and border zone of 11βHSD1i (n=8-9) and vehicle (n=6-8). Collagen organisation assessed using polarised light on picrosirius red stained infarct (d, e) and percent collagen that appears red-orange or green-yellow in both groups (f). Relative expression of LOX in both treatment groups (g). Scale bar = 50 µm Data are presented as mean±SEM. ns: not significant, **P<0.01, *P<0.05, two-way ANOVA with Bonferroni’s post-hoc (c, f) or two-tailed, unpaired t-test (b, g).

### 3.6 Proteomics analysis highlights alterations in proteins involved in extracellular matrix processing following 11ßHSD1i intervention

To further understand the impact 11BHSD1i on infarct repair, LV infarct tissue underwent proteomic analysis by mass spectrometry. The mass spectrometry proteomics data have been deposited to the ProteomeXchange Consortium via the PRIDE partner repository with the dataset identifier PXD055401, these will be publicly available when the MS has been peer reviewed and accepted for publication. Initial analysis of output using the *Reactome* database highlighted the pathways most regulated by 11ßHSD1i treatment compared to vehicle (*Figure 8a*, and *Supplementary Table 5*). The top 10 pathways identified in *Reactome* included extracellular matrix organisation (7 proteins, including 4 in elastic fibre formation, 3 molecules associated with elastic fibres and 4 ECM proteoglycans). The top regulated pathways also included proteins involved in the immune system (5), in protein metabolism (5) and gene expression (1). Further analysis in the *Gene Ontology Network (GO)* confirmed enrichment of proteins involved in ECM organisation and the complement cascade (*Figure 8b*). Differential expression analysis in *limma* identified specific proteins in these 2 pathways that were regulated by 11BHSD1i at 28 days after MI (*Figure 9a* and *Supplementary Table 5-6*). Interestingly, this analysis also highlighted a significant reduction in B-type natriuretic peptide (BNP), a protein associated with increased wall stress, consistent with improved cardiac function following 11BHSD1i administration. Network analysis in the *STRING* database identified association between proteins regulated by 11BHSD1i treatment and again primarily highlights proteins involved in extracellular matrix organisation, and links of these to complement related proteins (*Figure 9b*).

**Figure 7:**
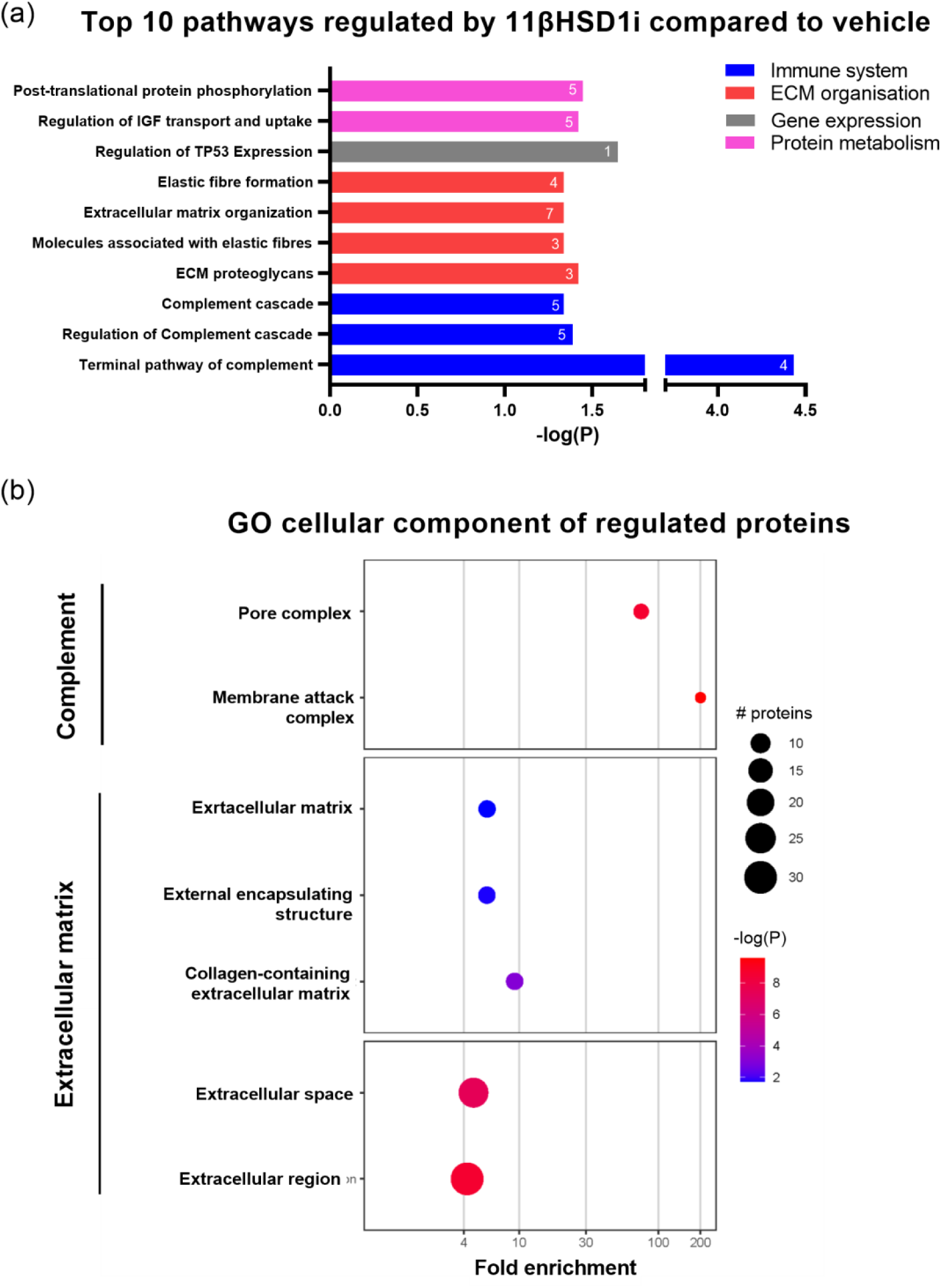
Extracellular matrix pathways and components are regulated by 11βHSD1i. Top ten regulated pathways analysed via Reactome in the peri-infarct of 11βHSD1i -treated pigs (n=9) compared to vehicle (n=9, a). Bar length represents P-value (longer, more significant). Numbers on bars refer to the number of regulated proteins involved in each pathway. Gene Ontology (GO) terms for cellular components of regulated proteins (b). Fold enrichment correlates with over-representation of each component. Bubble size corresponds to number of regulated proteins involved in each compartment and colour corresponds to P-value (blue=higher, red=lower). All P-value (a, b) are adjusted for multiple comparison and only P<0.05 enriched pathways or cellular compartments are represented.

**Figure 8.**
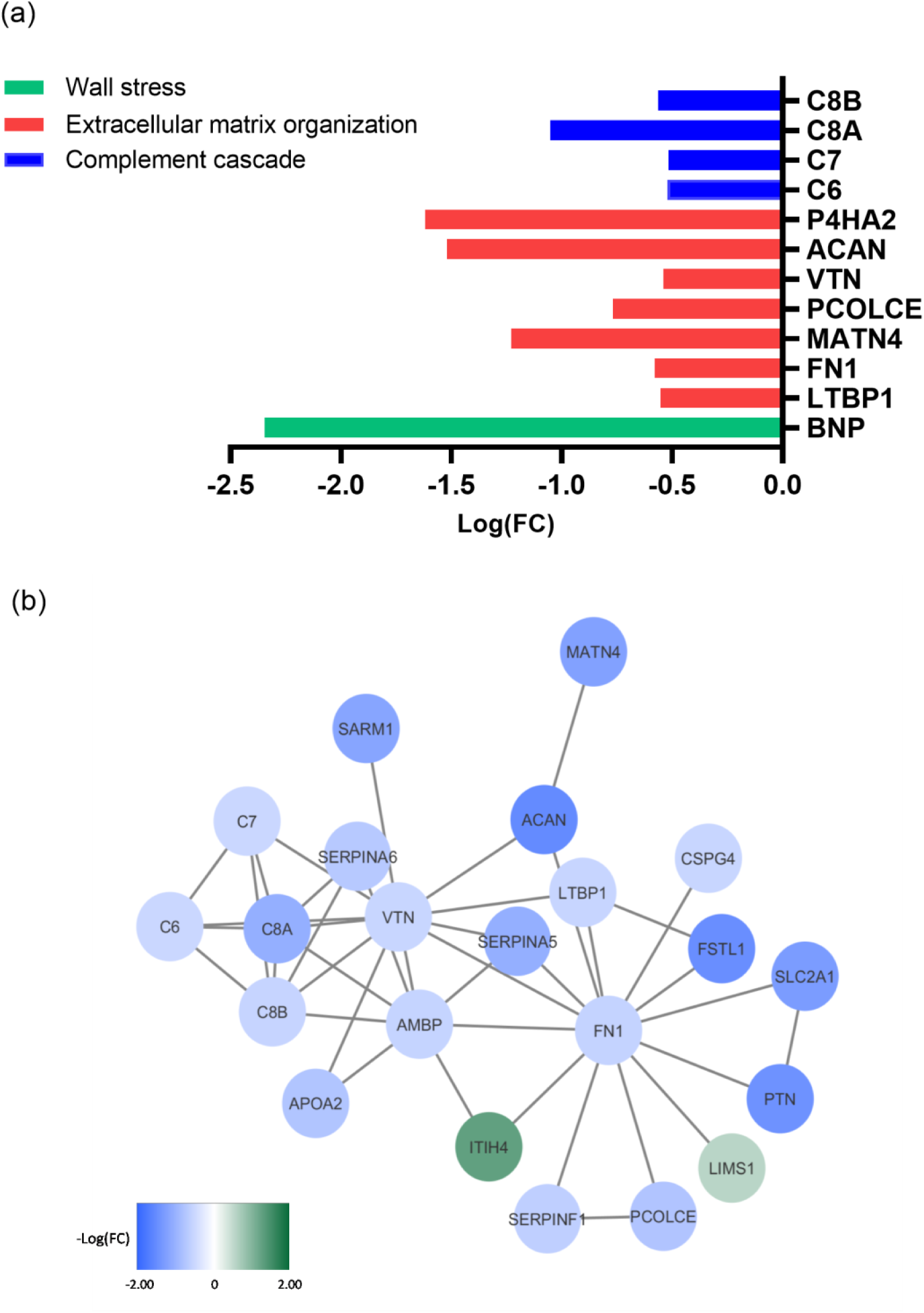
Proteins involved in extracellular matrix organisation, complement cascade and wall stress are downregulated by 11βHSD1i. Differential expression analysis of peri-infarct region (a). Bars correspond to fold change (FC) of proteins in 11βHSD1i (n=9) compared to vehicle (n=9). A network of the all differentially expressed proteins produced by STRING (b). Colour bar represents downregulated (blue) and upregulated (green) proteins in 11βHSD1i compared to vehicle, and shade corresponds to FC (darker=bigger change). All P-value (a, b) are adjusted for multiple comparison and only P<0.05 enriched pathways or cellular compartments are represented.

## 4. DISCUSSION

The primary aim of this study was to determine the potential for repurposing of pharmacological inhibitors of 11ßHSD1 to prevent adverse structural and functional cardiac remodelling following MI. The findings show that intervention for 4 weeks from 2 days after MI with the 11ßHSD1i AZD8293, in a translational percutaneous balloon mini-pig MI/reperfusion model treated with guideline recommended clinical therapies, successfully prevented functional decline and ventricular dilatation relative to vehicle treatment. Novel application of mass spectrometry imaging identified the repairing infarct as a key target for 11ßHSD1i post-MI, and proteomics revealed altered regulation of extracellular matrix processing in 11ßHSD1i treated pigs.

In the early period post-MI, HPA activation in response to a myocardial insult results in an acute rise in plasma GCs. Thereafter GCs are removed from the circulation by the action of the 11ßHSD2 in the kidney, generating the metabolite cortisone that is a substrate for tissue specific regeneration of active cortisol in cells expressing 11ßHSD1 (Chapman, Holmes & Seckl, 2013). In the heart *HSD11B1* is expressed in cardiomyocytes, and particularly highly in fibroblasts, as well as in neutrophils and macrophages recruited to the heart after MI (Gray, White, Castellan, McSweeney & Chapman, 2017). Cardiomyocyte damage in response to ischaemia quickly activates inflammatory cell recruitment to the heart after MI (Hilgendorf, Frantz & Frangogiannis, 2024). While this is essential for repair, over-recruitment is detrimental and can lead to cardiac rupture. Previously we showed that GCs regenerated in cardiac fibroblasts by 11ßHSD1 have an important role post-MI in suppressing the early recruitment of inflammatory cells by CXCL2 and CXCL4 (Mylonas et al., 2017). Regeneration in cardiomyocytes may also contribute to the early known cardioprotective actions of GCs in the heart (Libby, Maroko, Bloor, Sobel & Braunwald, 1973; Skyschally et al., 2004; Tokudome et al., 2009). In the present study pharmacological intervention with 11ßHSD1i was therefore begun from 2 days after MI to preserve any initial protective influence of 11ßHSD1 activity. Delaying intervention did not however prevent the longer-term benefits of 11ßHSD1i in terms of preservation of cardiac function and prevention of ventricular dilatation. Ejection fraction was improved by more than a clinically significant 5% in 11ßHSD1i-treated pigs.

In our previous mouse studies, improved function in *Hsd11b1*KO post-MI followed promotion of alternative macrophage polarisation, neovascularisation, vessel maturation and prevention of infarct expansion during repair (McSweeney et al., 2010; White et al., 2016). In the present study, in a more translational mini-pig model, there was clear evidence for ongoing infarct repair at study termination 28 days after MI, with the presence of FAP^+ve^ activated fibroblasts and pro-repair CD163^+ve^ macrophages (Schnitter et al., 2024). Application of mass spectrometry imaging (Israr, Bernieh, Salzano, Cassambai, Yazaki & Suzuki, 2020) to pig heart tissue sections collected after *in vivo* AZD8329 administration showed clear co-localisation of the 11ßHSD1i with the repairing tissue, supporting a role for 11ßHSD1 in this area. Identification by proteomics of modification in complement pathways in infarct tissue from 11ßHSDi treated pigs is consistent with an impact on the inflammatory response. However, neither macrophage density nor the extent of neovascularisation or infarct size were influenced by inhibition of 11ßHSD1. We cannot rule out an influence of the concurrent drug intervention in both groups that may have masked any effect here as both statins and ACE inhibitors can influence inflammation and angiogenesis (Bryniarski, Nazimek & Marcinkiewicz, 2022; Li, Zhao, Zhou, Chen & Li, 2004; Martovytskyi, Shelest & Kravchun, 2020; Weis, Heeschen, Glassford & Cooke, 2002). The timing of intervention, and the use of female pigs in a reperfusion model, rather than permanent coronary ligation in male mice, are also distinct from previous studies. Neovascularisation in the infarct border is less prominent in the pig heart repair process (Gallet et al., 2017; Liao et al., 2021; Rios-Navarro et al., 2018), which, more akin to human, tends instead to show evidence of collateral vessel expansion between the infarct and healthy myocardium (Schwartz et al., 1984). The mechanism underlying preserved structure and function in the translational pig model with 11ßHSD1i appears to be distinct from that previously reported in the mouse.

Change in scar structure can also impact on remote ventricular remodelling, and the reduction in the extent of BNP in the infarct border here is consistent with reduced wall stress after 11ßHSD1i. While there was no change in the amount of collagen or expression of collagen isoforms, birefringence imaging did support a potential change in collagen processing. This was accompanied by a reduction in expression of *LOX*, encoding the collagen cross linking enzyme lysyl oxidase, suppression of which has previously been linked to improved functional outcomes after MI (Gonzalez-Santamaria et al., 2016; Ma et al., 2023; Vukicevic et al., 2022). Proteomic analysis also pointed to extracellular matrix organisation as a major target of 11ßHSD1i, with reductions in fibronectin, procollagen C-endopeptidase enhancer (PCOLCE) and prolyl 4-hydroxylase subunit alpha 2 (P4HA2) proteins involved in collagen processing. Suppression of fibronectin was previously observed in mouse cardiac fibroblasts after *Hsd11b1* knockdown (Wang et al., 2022) and ejection fraction was improved post-MI by fibronectin inhibition and by fibroblast specific knockout (Valiente-Alandi et al., 2018). PCOLCE is also targeted by mineralocorticoid receptor antagonists that improve cardiac function after MI in mice (Kessler-Icekson, Schlesinger, Freimann & Kessler, 2006).

In the heart GCs regenerated by 11ßHSD1 can activate both glucocorticoid receptors and mineralocorticoid receptors (Gray, White, Castellan, McSweeney & Chapman, 2017), both implicated in repair after MI (Fraccarollo et al., 2008; Galuppo et al., 2017; Mihailidou, Loan Le, Mardini & Funder, 2009; Mylonas et al., 2017). 11ßHSD1 additionally regulates tissue availability of biologically active oxysterols, through the interconversion of 7 keto and 7 beta hydroxycholesterol (Mitic et al., 2013). The profile of 11ßHSD1i is therefore distinct from that of mineralocorticoid antagonists, currently recommended to prevent development of HF post-MI (McDonagh et al., 2021), and do not cause hyperkalaemia, that can limit the clinical use of this approach (Vizzardi et al., 2014).

11ßHSD1i are safe in humans and have beneficial metabolic effects (Anderson & Walker, 2013; Villemagne et al., 2024; Webster et al., 2017; Wilton-Waddell et al., 2024). The present translational study shows that administration after MI results in a clinically significant improvement in ejection fraction, even with concurrent relevant therapy. Together with evidence that tissue expression of *HSD11B1* is increased in ageing (Tiganescu et al., 2013) when the risk of MI is higher, this study provides strong support for the repurposing of 11ßHSD1i in the post-MI setting to prevent the development of HF.

## Abbreviations

αSMA: alpha smooth muscle actin
αSMA ACE: angiotensin converting enzyme
ARNI: Angiotensin Receptor Neprilysin Inhibitor
11ßHSD1: 11ß Hydroxysteroid Dehydrogenase type 1
11ßHSD1i: inhibitor of 11ß Hydroxysteroid Dehydrogenase type 1
ECM: extracellular matrix
FAP: fibroblast activation protein
FN: fibronectin
GC: glucocorticoid
Gd MRI: gadolinium enhanced magnetic resonance imaging
GR: glucocorticoid receptor
HF: heart failure
*Hsd11b1*: mouse 11ßHSD1 gene
*HSD11B1*: human 11ßHSD1 gene
IHD: ischaemic heart disease
LC-MS/MS: liquid chromatography mass spectrometry
LOX: lysyl oxidase
MALDI MSI: Matrix Assisted Laser Desorption-Ionisation Mass Spectrometry Imaging
MR: mineralocorticoid receptor
MRI: magnetic resonance imaging
PCOLCE: procollagen C-endopeptidase enhancer
P4HA2: prolyl 4-hydroxylase subunit alpha 2
SGLT2 inhibitor: inhibitor of sodium-glucose cotransporter 2
VT/VF: ventricular tachycardia/ventricular fibrillation
WGA: wheat germ agglutinin

## Acknowledgments

This study was supported by a Translational Grant from the British Heart Foundation (BHF TG/18/1/33048), a Novel and Emerging Technologies Grant from Heart Research UK (RG2694), by a Wellcome Trust (One Health Models of Disease) studentship to SAD (218471/Z/19/Z) and by a Wellcome Trust Multiuser Equipment Grant, (208402/Z/17/Z). NLM is supported by a Chair Award from the British Heart Foundation (CH/F/21/90010). AZ provided AZD8293 and supporting material through their Open Innovation Programme. The authors gratefully acknowledge the University of Bristol Translational Biomedical Research Facility, the University of Edinburgh Clinical Research Imaging Centre (CRIC), the Mass Spectrometry Core of the Edinburgh Clinical Research Facility, the Institute of Genetics and Cancer Mass Spectrometry Facility, Bioquarter SuRF Histology Facility and the Scottish Instrumentation and Resource Centre for Advanced Mass Spectrometry (SIRCAMS) for their expertise & assistance in this work.

## Conflict of interest statement

BRW and SPW are inventors on patents relating to 11ßHSD1 inhibitors which are owned by the University of Edinburgh and licensed to Actinogen Biomedical which is developing an 11ßHSD1 inhibitor for use in brain disorders. BRW is also a consultant for Actinogen Biomedical. AF is a full-time employee of AZ and may hold company shares and AW was a full-time employee of AZ at the time of the research and may hold company shares. None of the other authors have declared a conflict of interest.

