## Supplemental Material for "Pharmacological inhibition of 11ßhydroxysteroid dehydrogenase type 1 after myocardial infarction targets extracellular matrix processing and preserves cardiac function in a translational mini-pig model"

1. **Supplementary Figures**

**Supplementary Figure 1** Study Protocol: Adult female Goettingen mini-pigs received aspirin and simvastatin orally from 5 days prior to surgery for induction of MI by temporary coronary artery occlusion by balloon inflation. Amiodarone and heparin were infused intravenously during surgery only. Oral ramipril and and clopidogrel were added from 1 day after surgery. 11ßHSD1inhibitor (11ßHSD1i) or vehicle were administered orally from 2 days after MI induction. All oral treatment was continued until 1 day prior to study completion 28 days after MI. MRI for function (cine) and infarct size (LGE) were completed at 1 day and 28 days after MI.

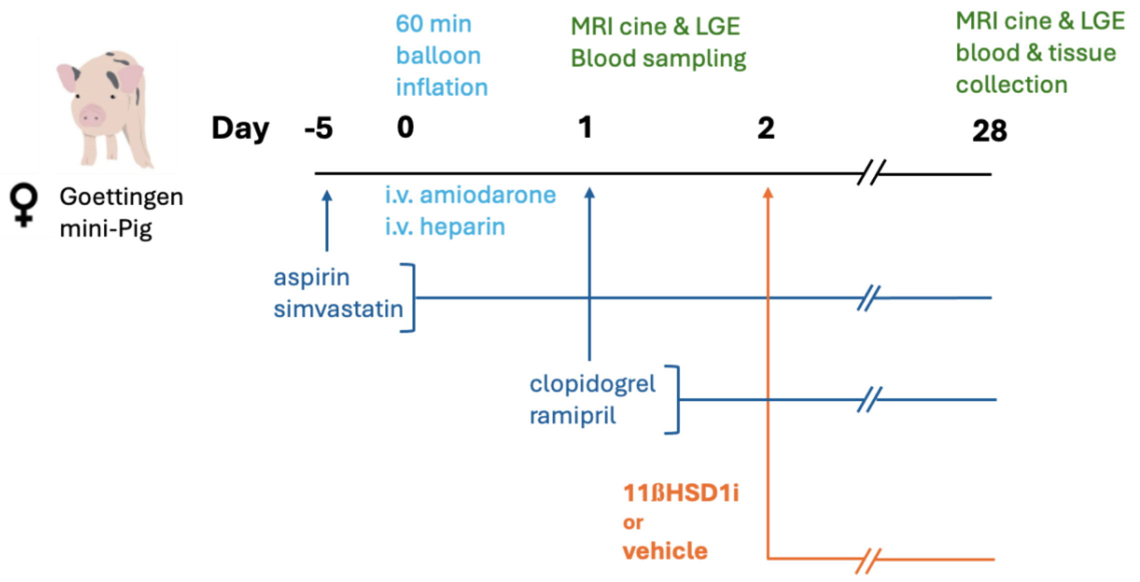

**Supplementary Figure 2** CD163$^{+ve}$ cell density in the infarct area of vehicle and standard 11βHSD1i treated pigs 28d post-MI. Every measurement (dot) represents the density of CD163$^{+ve}$ cells measured in 20 random boxes (1.6mm$^2$) in the infarct of each animal. Data presented as mean ± SEM. ns: not significant, unpaired t-test.

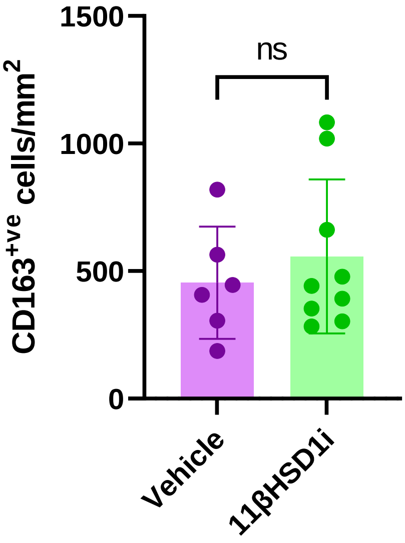

### 1. Supplementary Tables

**Supplementary Table 1:** List of TaqMan Gene Expression assays used in this study

| **Gene** | **Gene name** | **Species** | **TaqMan Assay ID** |
|---|---|---|---|
| COL1A1 | Collagen Type I | Pig | Ss03373340_m1 |
| COL3A1 | Collagen Type III | Pig | Ss04323794_m1 |
| FN1 | Fibronectin | Pig | Ss03373673_m1 |
| HMBS | Hydroxymethylbilane synthase | Pig | Ss03388782_g1 |
| HPRT1 | Hypoxanthine Phosphoribosyltransferase 1 | Pig | Ss03388274_m1 |
| LOX | Lysyl oxidase | Pig | Ss04323105_m1 |
| TBP | TATA-Box Binding Protein | Human | Hs00427620_m1 |

**Supplementary Table 2** qPCR reaction mix components

| **Component** | **Volume per reaction (µL)** |
|---|---|
| TaqMan Fast Advanced Master Mix (2X) | 5 |
| TaqMan Assay | 0.5 |
| Nuclease-free water | 3.5 |
| Total reaction mix | 9 |

**Supplementary Table 3** qPCR cycles program

| **Stage** | **Number of cycles** | **Step** | **Temperature (C)** | **Time** | **Ramp rate (C/s)** |
|---|---|---|---|---|---|
| Preincubation | 1 | 1 | 95 | 5 min | 4.8 |
| Amplification | 50 | 1 | 95 | 10 sec | 4.8 |
| | | 2 | 60 | 30 sec | 2.5 |
| Cooling | 1 | 1 | 40 | 30 sec | 2.5 |

**Supplementary Table 4** Functional parameters during MRI at the acute and chronic timepoints after MI in vehicle and 11βHSD1i treated pigs.

| **Parameter** | **Acute** | | **P-value** | **Chronic** | | **P-Value** |
|---|---|---|---|---|---|---|
| | Standard therapy (n=9) | +11βHSD1i (n=11) | | Standard therapy (n=9) | +11βHSD1i (n=11) | |
| **Heart rate (beat/min)** | 127 ± 8 | 115 ± 5 | 0.2035 | 115 ± 10 | 116 ± 5 | 0.9408 |
| **Stroke volume (ml)** | 18.30 ± 1.99 | 18.31 ± 1.10 | 0.9981 | 18.48 ± 1.74 | 20.52 ± 1.59 | 0.2625 |
| **Cardiac output (L/min)** | 2.40 ± 0.25 | 2.08 ± 0.12 | 0.3543 | 2.05 ± 0.11 | 2.35 ± 0.08 | *0.0399* |
| **Cardiac index (L/min/m$^2$)** | 2.3 ± 0.24 | 2.10 ± 0.10 | 0.5427 | 2.59 ± 1.78 | 3.10 ± 0.97 | *0.0163* |

| | | | | | | |
|---|---|---|---|---|---|---|
| **Myocardial mass (g)** | 28.40 ± 1.38 | 30.63 ± 1.212 | 0.2426 | 31.94 ± 1.18 | 31.93 ± 1.70 | 0.9952 |

**Supplementary Table 5** Reactome analysis tool results table, including all significantly enriched pathways. Disease-specific pathways were excluded. Rxn: reaction. FDR: false-discovery rate

| Pathway ID | Pathway name | Entities found | Entities total | Entities ratio | Entities P-value | Entities FDR | Rxn found | Rxn total | Rxn ratio | Species name | Submitted entities found |
|---|---|---|---|---|---|---|---|---|---|---|---|
| R-HSA-166665 | Terminal pathway of complement | 4 | 8 | 5.14E-04 | 1.12E-07 | 3.68E-05 | 5 | 5 | 3.36E-04 | Homo sapiens | C6 C7 C8B C8A |
| R-HSA-6804754 | Regulation of TP53 Expression | 2 | 4 | 2.57E-04 | 2.11E-04 | 0.022704768 | 4 | 5 | 3.36E-04 | Homo sapiens | PRDM1 |
| R-HSA-8957275 | Post-translational protein phosphorylation | 5 | 109 | 0.007000193 | 2.77E-04 | 0.022704768 | 1 | 1 | 6.72E-05 | Homo Sapiens | DNAJC3 FN1 APOA2 LTBP1 FSTL1 |
| R-HSA-3000178 | ECM proteoglycans | 4 | 79 | 0.005073534 | 8.08E-04 | 0.038059505 | 8 | 23 | 0.001544868 | Homo Sapiens | ACAN VTN FN1 MATN4 |
| R-HSA-977606 | Regulation of Complement cascade | 5 | 139 | 0.008926851 | 8.27E-04 | 0.038059505 | 4 | 42 | 0.002821064 | Homo Sapiens | VTN C6 C7 C8B C8A |
| R-HSA-2129379 | Molecules associated with elastic fibres | 3 | 37 | 0.002376212 | 9.97E-04 | 0.040864466 | 5 | 10 | 6.72E-04 | Homo sapiens | VTN FN1 LTBP1 |
| R-HSA-166658 | Complement cascade | 5 | 156 | 0.010018624 | 0.001375744 | 0.046303127 | 9 | 72 | 0.00483611 | Homo sapiens | VTN C6 C7 C8B C8A |
| R-HSA-1474244 | Extracellular matrix organization | 7 | 328 | 0.0210648 | 0.001641924 | 0.046303127 | 34 | 319 | 0.021426652 | Homo sapiens | ACAN VTN P4HA2 FN1 PCOLCE MATN4 LTBP1 |
| R-HSA-1566948 | Elastic fibre formation | 3 | 45 | 0.002889988 | 0.001740407 | 0.046303127 | 5 | 17 | 0.001141859 | Homo sapiens | VTN FN1 LTBP1 |
| R-HSA-9700645 | ALK mutants bind TKIs | 2 | 12 | 7.71E-04 | 0.001847936 | 0.046303127 | 1 | 1 | 6.72E-05 | Homo sapiens | EML4 FN1 |
| R-HSA-201556 | Signaling by ALK | 3 | 46 | 0.00295421 | 0.001852125 | 0.046303127 | 26 | 42 | 0.002821064 | Homo sapiens | PRDM1 PTN |

**Supplementary Table 6** Differentially expressed proteins in 11βHSD1i compared to vehicle as analysed by *limma*. FC: fold change, p.mod: adjusted P-value for multiple testing

| Symbol | -logFC | p.mod |
|---|---|---|
| A0A075B7I5 | -1.69844 | 0.041689 |
| A0A075B7I6 | -0.69872 | 0.104567 |
| SEPTIN5 | -1.20515 | 0.091142 |
| NRBP1 | -0.57256 | 0.249245 |
| EMC4 | 0.937203 | 0.293419 |
| DLGAP4 | -0.60505 | 0.12274 |
| ARFGAP1 | 0.862995 | 0.56273 |
| GRIPAP1 | -0.55202 | 0.378611 |
| LOC100514282 | -0.51172 | 0.333412 |
| EIF2B1 | -0.74271 | 0.150065 |
| LIMS2 | -0.58711 | 0.322272 |
| GPNMB | -1.05292 | 0.300661 |
| KIF2A | 0.75355 | 0.456443 |
| FAM120B | 0.554401 | 0.346292 |
| INPP5F | -0.82313 | 0.170899 |
| TRAPPC4 | 0.579382 | 0.539376 |
| PPP6R2 | -0.61381 | 0.112835 |
| COG6 | -1.15486 | 0.151154 |
| FBLN1 | 0.534601 | 0.469062 |
| A0A286ZLN6;A0A286ZUL8;A0A287A3T6;A0A287AUR4;A0A287AWQ8 | -0.59136 | 0.089429 |
| CARM1 | 0.757734 | 0.059827 |
| TPD52 | -1.96814 | 0.062367 |
| NALCN | -0.65826 | 0.644848 |
| DOT1L | 1.118722 | 0.522795 |
| LOC100739101;LOC102167481 | 0.700436 | 0.130507 |
| TCEA3 | 0.626072 | 0.387212 |
| ARHGEF1 | 0.580197 | 0.332753 |
| DIS3L2 | 0.689598 | 0.278783 |
| NUBP2 | -0.8412 | 0.245245 |
| D2HGDH | 0.512246 | 0.328551 |
| ABR | -0.53415 | 0.302749 |
| DAP | -0.58665 | 0.268501 |
| A0A286ZQQ8;A0A287AKW2;A0A5G2R1F2;I3LBK0 | -0.91659 | 0.47501 |
| SLAIN2 | 0.52869 | 0.386993 |
| NUP214 | -0.68429 | 0.348696 |
| RCSD1 | -0.67214 | 0.15618 |
| C1QC | -0.58287 | 0.167748 |
| SDF4 | -1.17076 | 0.035433 |
| A0A286ZSX6;A0A8W4F9N8 | -0.97281 | 0.252738 |
| OSBPL9 | -0.97064 | 0.360807 |
| HSDL1 | 0.83076 | 0.322395 |
| POLR1C | 0.799321 | 0.088458 |
| ITIH4 | 0.684761 | 0.604803 |

| | | |
|---|---|---|
| NAAA | 1.076367 | 0.13963 |
| TXLNA | -0.50674 | 0.138397 |
| RGS6 | -0.65463 | 0.333837 |
| NUP98 | 0.522149 | 0.597294 |
| A0A286ZUL8 | 2.64266 | 0.077902 |
| GNS | 0.878972 | 0.189232 |
| CCDC50 | -0.53221 | 0.178567 |
| RER1 | -2.39942 | 0.059619 |
| LOC106504547 | 0.762582 | 0.063712 |
| NLRP12L | -0.72223 | 0.376163 |
| DYNC1I1 | 0.911093 | 0.2397 |
| TRAK1 | -0.87226 | 0.327927 |
| NT5C3B | -0.54049 | 0.143486 |
| LTBP1 | -0.5526 | 0.021993 |
| COG5 | -0.86573 | 0.455724 |
| CLPTM1 | -0.52738 | 0.615844 |
| OLFML1 | -0.58106 | 0.236871 |
| EML3 | -0.90448 | 0.161541 |
| DTYMK | -0.54317 | 0.397694 |
| ERLEC1 | -1.29118 | 0.176629 |
| PLEKHA5 | -0.95123 | 0.199642 |
| BRD4 | 0.525755 | 0.490092 |
| AEBP1 | -0.81366 | 0.386379 |
| SRP72 | -0.54668 | 0.392584 |
| DBN1 | -0.75532 | 0.180663 |
| PYCR1;PYCR2 | 0.660223 | 0.301682 |
| CYP2U1 | 0.569894 | 0.337988 |
| GFAP | -0.7925 | 0.412148 |
| STX8 | -0.55905 | 0.314855 |
| GPX7 | -0.68355 | 0.49218 |
| CLNS1A | 0.526838 | 0.286084 |
| TLE5 | 1.325409 | 0.018775 |
| RASA1 | -1.25508 | 0.117977 |
| PYCR3 | -0.99246 | 0.120403 |
| PLG | -0.73995 | 0.449301 |
| BORCS5 | -1.06454 | 0.351053 |
| TATDN3 | 0.99857 | 0.197023 |
| TMEM30A | -0.60172 | 0.208511 |
| PTGES3L | 0.717392 | 0.318909 |
| ASH2L | 0.686163 | 0.360145 |
| DAAM1 | 0.771852 | 0.41069 |
| RRM1 | -1.54436 | 0.139593 |
| NOA1 | 1.001063 | 0.146112 |
| CES1 | -1.04653 | 0.359698 |
| DAB2 | -0.63434 | 0.447988 |
| CLIC3 | -1.96751 | 0.037135 |

| | | |
|---|---|---|
| AP3S2 | -0.60817 | 0.307034 |
| DYNLT1 | -0.60579 | 0.322829 |
| VPS16 | -0.51725 | 0.372991 |
| GRK2 | 0.564712 | 0.275291 |
| MFAP2 | -0.79431 | 0.18483 |
| BROX | 1.344043 | 0.053012 |
| CC2D1A | 0.767449 | 0.426412 |
| ISG15 | -0.59284 | 0.582272 |
| TANC1 | -1.80298 | 0.024625 |
| PLPP3 | -1.13334 | 0.31773 |
| CCDC9 | -0.614 | 0.04171 |
| EEFSEC | -1.98726 | 0.031036 |
| TBC1D15 | -0.65148 | 0.21687 |
| LMNA | -0.86751 | 0.683094 |
| KPNA6 | -0.83083 | 0.258109 |
| ABHD4 | -1.02677 | 0.12299 |
| FAM114A1 | -1.01938 | 0.048149 |
| RABEP2 | -1.36089 | 0.020113 |
| RAB8B | 1.81956 | 0.058583 |
| LOC106504547 | 0.81482 | 0.09076 |
| CSAD | -0.86506 | 0.155309 |
| EML4 | -1.37039 | 0.001943 |
| ACAT2 | -0.92476 | 0.05674 |
| CNBP | -1.77483 | 0.050473 |
| BICD2 | -0.67805 | 0.364403 |
| ATG7 | -0.66393 | 0.133016 |
| TRIOBP | -0.63281 | 0.195336 |
| PDS5B | 0.53871 | 0.428494 |
| DDO | -1.57814 | 0.069219 |
| EIF2B4 | -0.62564 | 0.320593 |
| FBLN1 | -0.54054 | 0.127875 |
| STAB1 | -0.54203 | 0.218213 |
| PLOD2 | -0.70155 | 0.307324 |
| MYH8 | 0.592887 | 0.450513 |
| RCC2 | -0.95606 | 0.137652 |
| A0A287AP28 | -1.73505 | 0.019612 |
| EPB41L3 | -0.66581 | 0.377865 |
| KRT75 | -0.62145 | 0.603484 |
| RAP1B | 0.761063 | 0.259445 |
| MTHFSD | 0.519256 | 0.329261 |
| TMEM186 | -0.96136 | 0.373231 |
| A0A287ARL5;A0A287B5G0 | -1.54135 | 0.117891 |
| QRSL1 | 0.752256 | 0.171305 |
| C8B | -0.56232 | 0.013532 |
| KPNA4 | -0.80042 | 0.204424 |
| PLVAP | -0.61718 | 0.217791 |

| | | |
|---|---|---|
| DMTN | 0.656027 | 0.298872 |
| A0A287AUR4 | -0.71234 | 0.58685 |
| ILKAP | 0.602248 | 0.135664 |
| RPL37 | -0.50148 | 0.113253 |
| MATN2 | -0.60614 | 0.284975 |
| CAPZB | 1.176209 | 0.13079 |
| PPT1 | -0.51524 | 0.495316 |
| SERPINF1 | -0.63147 | 0.034668 |
| PKN1 | 0.678431 | 0.207782 |
| SERPINE2 | -0.97071 | 0.220378 |
| CAMK1 | -1.37995 | 0.143796 |
| CA14 | 0.819709 | 0.260184 |
| NANS | -0.70083 | 0.289019 |
| PRDM1 | 1.189191 | 0.020926 |
| STRBP | -1.0721 | 0.412946 |
| ATG5 | -1.57641 | 0.024661 |
| ATP13A1 | -1.2488 | 0.079654 |
| HOMER2 | -0.62973 | 0.307144 |
| NEK3 | -0.72008 | 0.099218 |
| RHOG | 0.669936 | 0.213073 |
| PUS1 | 0.810992 | 0.047045 |
| CCDC166 | -1.04541 | 0.313444 |
| RAB4B | -1.42084 | 0.014948 |
| SRR | -0.77278 | 0.286713 |
| NARS2 | -0.64046 | 0.459333 |
| CA2 | -0.7577 | 0.137255 |
| ACAD10 | 1.740036 | 0.130911 |
| RARRES2 | -0.78992 | 0.35537 |
| ACAN | -1.76946 | 0.041534 |
| PTMS | -0.6827 | 0.150356 |
| CAPG | -0.70838 | 0.160628 |
| CPQ | -0.58395 | 0.569597 |
| DDAH1 | -0.89055 | 0.191871 |
| TMEM205 | -0.78151 | 0.239889 |
| POFUT2 | -0.51734 | 0.348891 |
| SMARCB1 | 2.026292 | 0.061695 |
| PARS2 | 1.011432 | 0.047251 |
| PACS2 | -0.7434 | 0.241256 |
| DDRGK1 | -0.53618 | 0.157467 |
| MRPL51 | -1.10615 | 0.149024 |
| PDLIM7 | -0.62625 | 0.323466 |
| MSH4 | -0.8505 | 0.284091 |
| ITIH4 | 0.635329 | 0.71755 |
| HBS1L | -0.65496 | 0.133793 |
| FGF2 | -0.82615 | 0.12865 |
| COX6A1 | -0.60166 | 0.399858 |

| | | |
|---|---|---|
| MRE11 | 1.812298 | 0.029734 |
| HMGB3 | -2.13516 | 0.040507 |
| KRT2 | 0.946779 | 0.303926 |
| LOC110259265 | -1.12604 | 0.12963 |
| AP2S1 | -0.69507 | 0.205769 |
| DCAF7 | -0.83757 | 0.362098 |
| EFEMP2 | -0.90763 | 0.413083 |
| ACSL4 | -1.23969 | 0.101098 |
| FGL1 | 0.690843 | 0.420023 |
| FSTL1 | -1.4741 | 0.036613 |
| TXNL1 | -0.7872 | 0.391812 |
| RPS6KA5 | 0.798769 | 0.159333 |
| CNPY4 | -1.94472 | 0.036998 |
| EEPD1 | 0.618021 | 0.43029 |
| B3GAT3 | -0.61896 | 0.216481 |
| MARCKS | -0.70366 | 0.00296 |
| TM9SF4 | -1.23589 | 0.278834 |
| GBP1 | 0.612341 | 0.101038 |
| LAD1 | 1.341043 | 0.060349 |
| SRSF4 | 0.687082 | 0.498005 |
| P3H1 | -0.87348 | 0.330049 |
| GOLGA3 | -1.5908 | 0.093402 |
| LYZ | 0.796032 | 0.365792 |
| FKBP10 | -0.68654 | 0.118317 |
| COL14A1 | -0.53922 | 0.109333 |
| SCN3B | -0.99765 | 0.225992 |
| SERPINA5 | -1.04866 | 0.011639 |
| NUP35 | -0.76999 | 0.207842 |
| PDLIM3 | -1.18949 | 0.130905 |
| LHPP | -0.58825 | 0.312963 |
| CTPS2 | -1.23756 | 0.282445 |
| TARS2 | -0.70359 | 0.097677 |
| PAK4 | -2.47273 | 0.006786 |
| PTER | -0.69982 | 0.297596 |
| RSL1D1 | -0.50218 | 0.432776 |
| APOA2 | -0.76373 | 0.000857 |
| COPZ2 | -1.14395 | 0.078897 |
| AP4E1 | -0.69444 | 0.205645 |
| SCFD2 | -1.20391 | 0.13441 |
| CD109 | -1.25793 | 0.094185 |
| N6AMT1 | -0.76994 | 0.111743 |
| C6orf136 | 0.580077 | 0.135347 |
| PRRC1 | -1.67772 | 0.100219 |
| PARP4 | -0.72348 | 0.428384 |
| ODR4 | -0.90042 | 0.116701 |
| PFDN4 | -3.08625 | 0.041652 |

| | | |
|---|---|---|
| FAP | 1.133533 | 0.255702 |
| DRAP1 | -0.68632 | 0.426038 |
| SNW1 | -0.68068 | 0.242146 |
| ITIH4 | 1.230023 | 0.009133 |
| FN1 | -0.57808 | 0.035123 |
| PNKP | -0.59736 | 0.322656 |
| TPK1 | 0.71305 | 0.386947 |
| COL11A1 | -0.90921 | 0.074354 |
| MCM5 | 0.593588 | 0.411195 |
| RSC1A1 | 0.65027 | 0.409655 |
| AGPAT1 | 0.655436 | 0.156222 |
| ANKRD2 | -1.48126 | 0.163053 |
| A0A5G2QM05 | -0.72291 | 0.025878 |
| ABCF3 | -0.52611 | 0.464129 |
| ARIH1 | -1.21582 | 0.071092 |
| CYB5R3 | -2.49035 | 0.048487 |
| CPLX1 | 0.626964 | 0.072179 |
| FARP1 | -1.64126 | 0.111921 |
| LIMS1 | 0.55178 | 0.010591 |
| SDC2 | -0.58573 | 0.180639 |
| BCL2L13 | -0.86922 | 0.305731 |
| PPIH | -0.60799 | 0.308095 |
| EIF3M | -1.72803 | 0.151789 |
| DIS3 | 0.773609 | 0.235818 |
| PFN2 | 0.86646 | 0.501893 |
| DDX46 | -0.56366 | 0.196445 |
| KRT3 | 0.912471 | 0.093123 |
| RAB11FIP5 | -0.73848 | 0.345318 |
| CA4 | 0.680275 | 0.016505 |
| RBP1 | -0.51822 | 0.386168 |
| PSAT1 | -0.576 | 0.313135 |
| CTHRC1 | -1.66716 | 0.151717 |
| LOC100519082;LOC100626247 | 0.960895 | 0.16095 |
| LOC100158003 | 1.369261 | 0.414379 |
| SMARCA4 | -0.5212 | 0.33657 |
| TPM4 | -0.84295 | 0.084787 |
| STX16 | -1.21276 | 0.079277 |
| A0A5G2QXT5 | -1.00176 | 0.064019 |
| SLC30A9 | -0.58539 | 0.444489 |
| DNAJC3 | 1.852179 | 0.017611 |
| ELAC2 | -0.84212 | 0.185847 |
| MRPS18A | 1.101515 | 0.177191 |
| RAC2 | -1.20178 | 0.135321 |
| FAM107B | 0.736117 | 0.01188 |
| ALDH1A2 | -2.21539 | 0.018667 |
| COLEC12 | -0.77519 | 0.228347 |

| | | |
|---|---|---|
| ROR1 | -1.02286 | 0.098661 |
| MCAM | -0.65477 | 0.306559 |
| TBCE | -0.72053 | 0.029722 |
| GNG10 | -1.81653 | 0.184924 |
| MMP2 | -0.84226 | 0.31061 |
| FKBP11 | -2.25391 | 0.033371 |
| PPP3CB | -1.1292 | 0.107909 |
| CORO7 | 0.68368 | 0.450623 |
| C6 | -0.52106 | 0.029408 |
| SMOC1 | -1.07197 | 0.035734 |
| ENOSF1 | -0.60194 | 0.099255 |
| PDIA5 | -0.63839 | 0.303329 |
| SCAMP4 | -1.29961 | 0.052242 |
| PON2 | 0.766365 | 0.173111 |
| AS3MT | -1.34917 | 0.053206 |
| ATP6AP1 | -0.82603 | 0.093044 |
| TUBB3 | 1.217201 | 0.201298 |
| SULT1C4 | -0.75591 | 0.285673 |
| IFIT3 | 0.649508 | 0.445286 |
| BGN | -0.99434 | 0.242308 |
| SHC1 | 0.624557 | 0.100583 |
| PADI2 | -1.45927 | 0.033277 |
| EMC6 | -0.80028 | 0.25357 |
| CRTAP | -0.76777 | 0.262436 |
| A0A8W4F8D3 | -0.69774 | 0.008317 |
| ABCD1 | -0.54097 | 0.488791 |
| NEBL | 0.721769 | 0.343341 |
| NDUFC1 | 0.658991 | 0.318854 |
| FTL | -1.02935 | 0.169435 |
| A0A8W4FF43;A0A8W4FJA1 | -0.85557 | 0.151286 |
| DHRS7C | 0.661713 | 0.081197 |
| SUSD2 | -0.51589 | 0.53417 |
| FHL3 | -0.60863 | 0.2695 |
| SELENOM | 0.542175 | 0.53258 |
| GNAI3 | -0.73795 | 0.218102 |
| NBAS | 0.683751 | 0.266406 |
| P3H3 | -0.6458 | 0.515836 |
| COL1A2 | -0.53278 | 0.220089 |
| GBF1 | -0.53832 | 0.370782 |
| A0A8W4FPQ3 | -0.68757 | 0.419516 |
| SNRPE | -2.69343 | 0.049399 |
| COX8H | -1.50074 | 0.134985 |
| FYN | 1.057414 | 0.222401 |
| OLFML2A | 1.275515 | 0.073036 |
| PSMB8 | 0.620075 | 0.47749 |
| RAE1 | -0.5331 | 0.289359 |

| | | |
|---|---|---|
| DPM1 | -0.63151 | 0.217949 |
| SIGLEC1 | -0.87442 | 0.293027 |
| AK4 | -0.93855 | 0.219311 |
| CRABP1 | -0.90207 | 0.448723 |
| DYNLL1 | 1.086555 | 0.15902 |
| STAT6 | -1.24739 | 0.064742 |
| VPS37B | -1.61895 | 0.09573 |
| VCAN | -0.63126 | 0.093259 |
| RCN3 | -0.51592 | 0.270002 |
| DNAAF5 | -0.96449 | 0.185587 |
| LOC100523213 | -0.56818 | 0.036338 |
| FGF1 | 1.263067 | 0.3061 |
| MBLAC2 | -0.53184 | 0.531204 |
| GOLGA4 | -0.5139 | 0.29683 |
| MYL4 | -0.80294 | 0.011323 |
| MRC2 | -0.50039 | 0.247274 |
| PTRH1 | 0.54714 | 0.258299 |
| ARHGAP10 | -0.53961 | 0.467415 |
| ACTN3 | 1.692446 | 0.386697 |
| PNPO | -1.13193 | 0.169843 |
| ECI2 | 0.522372 | 0.021244 |
| FUNDC1 | -0.55486 | 0.122968 |
| FRZB | -1.67626 | 0.058388 |
| GATD1 | -0.60463 | 0.218426 |
| CHMP6 | -0.61868 | 0.284584 |
| RAB3GAP1 | -1.39108 | 0.049308 |
| SEC24D | 0.659632 | 0.354622 |
| CDK18 | 0.715077 | 0.226584 |
| LOC100524873 | 0.862711 | 0.181221 |
| LDHD | 0.857453 | 0.154855 |
| ABLIM1 | 0.622874 | 0.044747 |
| CILP2 | -1.17485 | 0.363307 |
| TEAD1 | 1.957303 | 0.017093 |
| TNN | -1.02238 | 0.088839 |
| C8A | -1.05196 | 0.024973 |
| MTRF1L | -0.59406 | 0.130991 |
| TMEM38A | 0.560512 | 0.45564 |
| EARS2 | -1.51845 | 0.040126 |
| PTGFRN | -0.64448 | 0.206243 |
| DNAH11 | 0.55833 | 0.559741 |
| PUM2 | 1.112518 | 0.0605 |
| UQCR11 | 0.523959 | 0.750329 |
| MATN4 | -1.22872 | 0.030295 |
| CHMP5 | -2.0062 | 0.067621 |
| S100A4 | -0.73424 | 0.291625 |
| PKP2 | 0.538998 | 0.099257 |

| | | |
|---|---|---|
| NAT10 | -0.60682 | 0.502437 |
| FZD7 | -0.59818 | 0.230486 |
| RPLP1 | 1.638082 | 0.311132 |
| GLYCTK | 1.466308 | 0.04151 |
| TEX264 | -0.66449 | 0.33747 |
| CSPG4 | -0.53516 | 0.023531 |
| PTCD2 | -0.53677 | 0.352612 |
| COL7A1 | -1.51691 | 0.120768 |
| MTREX | -0.84357 | 0.214166 |
| C9H11orf52 | -0.69771 | 0.574404 |
| GARIN1A | -0.53281 | 0.440686 |
| FBLN2 | -0.5165 | 0.124751 |
| CD63 | 0.53447 | 0.583557 |
| MST1 | -1.34209 | 0.016832 |
| KERA | -0.67469 | 0.502568 |
| ABCB6 | -1.24674 | 0.529507 |
| THBS1 | -0.52352 | 0.143167 |
| THTPA | 0.75283 | 0.416385 |
| SMPDL3B | -0.55312 | 0.331875 |
| DDX21 | -0.71307 | 0.349925 |
| COA4 | 0.801362 | 0.005845 |
| SLC25A5 | -1.16112 | 0.012473 |
| H2BC4 | -0.64874 | 0.278004 |
| RAB8A | -2.36975 | 0.011954 |
| RPL26L1 | -0.58232 | 0.355707 |
| SNRPG | -2.05947 | 0.137177 |
| SRSF11 | 0.701928 | 0.201389 |
| OLFML3 | -1.10176 | 0.195798 |
| SARM1 | -1.17786 | 0.048466 |
| P4HA2 | -1.61951 | 0.0435 |
| I3L6U3 | 1.20598 | 0.209146 |
| CCZ1 | -1.22924 | 0.147147 |
| LOC100737821 | 0.565097 | 0.247475 |
| LIPT2 | -0.73661 | 0.074608 |
| ARHGAP21 | 1.71365 | 0.073351 |
| PODN | -0.64318 | 0.215261 |
| PCOLCE | -0.76765 | 0.041717 |
| LOC110255237 | 1.36271 | 0.071401 |
| I3LFU3 | -0.64818 | 0.53354 |
| RPA3 | 0.793143 | 0.160849 |
| GDE1 | -0.98577 | 0.161271 |
| COA7 | -0.95642 | 0.1271 |
| KBTBD11 | -0.60726 | 0.433953 |
| OMA1 | 0.532728 | 0.434438 |
| IGF2BP2 | 0.808998 | 0.149439 |
| S100A2 | -1.91915 | 0.134039 |

| | | |
|---|---|---|
| MOGS | 0.544024 | 0.326747 |
| YY1 | -0.91953 | 0.139277 |
| TM9SF2 | -0.53418 | 0.426361 |
| CRP | 0.623321 | 0.239994 |
| CCN2 | -1.15067 | 0.063213 |
| TOP2A | -0.65512 | 0.286428 |
| CTSH | -0.68756 | 0.358657 |
| BCL2L1 | 0.717441 | 0.281006 |
| FABP4 | -0.57033 | 0.207588 |
| PDYN | -0.81917 | 0.386711 |
| AMBP | -0.56627 | 0.024253 |
| NPPB | -2.34525 | 0.050772 |
| UBC | 0.601729 | 0.083105 |
| NFI;NFIC | 0.653316 | 0.091224 |
| RNASE4 | -0.73457 | 0.162063 |
| P15980 | 0.857742 | 0.441436 |
| P15982 | 0.656981 | 0.482423 |
| IGFBP3 | -0.92662 | 0.090251 |
| APOA1 | -0.56969 | 0.051335 |
| FTL | -0.53339 | 0.281979 |
| SLC2A1 | -1.29517 | 0.048002 |
| NPPA | -0.64757 | 0.199994 |
| APOC3 | -0.53947 | 0.135954 |
| GUK1 | -1.12312 | 0.055355 |
| HSPA1A | 0.793332 | 0.198545 |
| VTN | -0.54096 | 0.006904 |
| NUDT2 | 0.508654 | 0.013705 |
| CFD | -1.24105 | 0.133752 |
| NPC1 | -0.755 | 0.444729 |
| ITIH4 | 1.241299 | 0.011933 |
| MYLK | -0.57846 | 0.670167 |
| PTN | -1.42472 | 0.011457 |
| HMGN2 | -0.57664 | 0.381256 |
| ACAN | -1.51995 | 0.043238 |
| ICA | -0.50064 | 0.013042 |
| OAS1 | 0.694577 | 0.575628 |
| PALM | 0.809687 | 0.247139 |
| GUSB | -0.50119 | 0.210016 |
| ENPP6 | -1.52671 | 0.152389 |
| CA3 | -0.93567 | 0.186061 |
| LIPE | 0.638135 | 0.366741 |
| ApoN | -0.56451 | 0.029637 |
| RETN | 0.748501 | 0.277071 |
| SPP2 | -1.28766 | 0.159593 |
| VARS2 | -0.89075 | 0.257673 |
| PPP1R14B | -0.61712 | 0.450894 |

| | | |
|---|---|---|
| MGP | -0.58763 | 0.180898 |
| OSTF1 | 1.028162 | 0.122346 |
| Serpina6 | -0.7071 | 0.003965 |
| BGN | -0.58344 | 0.219131 |
| SMAD4 | 1.490586 | 0.071851 |
| CHGB | 0.703042 | 0.276358 |
| F5 | -0.55948 | 0.208442 |
| MCT7 | 0.96851 | 0.404243 |
| SERPINA7 | 0.973544 | 0.176435 |
| FMOD | -1.22145 | 0.41807 |
| OLR1 | -0.67117 | 0.283417 |
| C7 | -0.51639 | 0.035489 |
